# Atomic Models of All Major Trans-Envelope Complexes Involved in Lipid Trafficking in *Escherichia Coli* Constructed Using a Combination of AlphaFold2, AF2Complex, and Membrane Morphing Simulations

**DOI:** 10.1101/2023.04.28.538765

**Authors:** Robert T. McDonnell, Nikhil Patel, Zachary J. Wehrspan, Adrian H. Elcock

## Abstract

In Gram-negative bacteria, several trans-envelope complexes (TECs) have been identified that span the periplasmic space in order to facilitate lipid transport between the inner- and outer-membranes. While partial or near-complete structures of some of these TECs have been solved by conventional experimental techniques, most remain incomplete. Here we describe how a combination of computational approaches, constrained by experimental data, can be used to build complete atomic models for four TECs implicated in lipid transport in *Escherichia coli*. We use DeepMind’s protein structure prediction algorithm, AlphaFold2, and a variant of it designed to predict protein complexes, AF2Complex, to predict the oligomeric states of key components of TECs and their likely interfaces with other components. After obtaining initial models of the complete TECs by superimposing predicted structures of subcomplexes, we use the membrane orientation prediction algorithm OPM to predict the likely orientations of the inner- and outer-membrane components in each TEC. Since, in all cases, the predicted membrane orientations in these initial models are tilted relative to each other, we devise a novel molecular mechanics-based strategy that we call “membrane morphing” that adjusts each TEC model until the two membranes are properly aligned with each other and separated by a distance consistent with estimates of the periplasmic width in *E. coli*. The study highlights the potential power of combining computational methods, operating within limits set by both experimental data and by cell physiology, for producing useable atomic structures of very large protein complexes.

## Introduction

Gram-negative bacteria, such as *Escherichia coli*, have cell envelopes composed of two lipid membranes that help both to confer antibiotic resistance (Lehman 2019) and to maintain the osmotic pressure of the cell (Rojas et al., 2014; Hwang et al., 2018). The two membranes, commonly referred to as the inner membrane (IM) and outer membrane (OM), are separated by an aqueous cellular space known as the periplasm (Mitchell 1961). Both membranes house a range of lipid types that may be asymmetrically compartmentalized to either the inner or outer leaflet of each membrane (Silhavy et al., 2010), with a key example being that the outer leaflet of the OM is primarily composed of lipopolysaccharide (LPS) (Silhavy et al., 2010). In order to maintain the integrity of both the IM and OM, bacteria have evolved protein-based systems that shuttle lipids across the periplasmic space; most of these are large, stable protein complexes that traverse the periplasm, and that are therefore termed trans-envelope complexes (TECs). Recent years have seen dramatic progress (reviewed in Giacometti et al., 2022) in elucidating the structures of TECs in *E. coli*, but due to their size and other experimental limitations, many of the available structures are missing known components.

In a parallel development, DeepMind’s deep learning-based prediction method, AlphaFold2, has revolutionized the field of computational structural biology, providing a way to routinely produce high-quality predictions of protein structures (Jumper et al., 2021). Trained on structures deposited in the RCSB (Berman et al., 2000), AlphaFold2 and related methods produce predictions typically within hours, thereby providing researchers with a powerful tool for acquiring useable structural information without the cost and time associated with experimental techniques. AlphaFold2 has already been shown to have a wide range of applications, such as identifying ligand binding sites (Wehrspan et al., 2021, Hekkelman et al., 2022), predicting or sampling alternative conformations (del Alamo et al., 2022; Stein & McHaourab, 2022), categorizing structural disorder (Wilson et al., 2022) and even increasing the resolution of experimental models (Terwilliger et al., 2022).

Following AlphaFold2’s release, considerable attention has been focused on adapting it, and related methods, to predict the structures of protein complexes. The DeepMind team has, for example, reported AlphaFold-Multimer (Evans et al., 2022) but similar competing methods have also been developed by other groups, such as RoseTTAfold (Baek et al., 2021), ColabFold (Mirdita et al., 2022), and AF2Complex (Gao et al., 2022). It is important to note that all these methods make internal use of neural network models based on the framework initially developed by DeepMind for AlphaFold2. Because AlphaFold2 produces high-quality structures for multidomain proteins across several model organisms (Tunyasuvunakool et al., 2021), and because the physical forces driving domain-domain interactions are largely the same as those driving protein-protein interactions (Keskin et al., 2008), many AlphaFold2-derived complex prediction approaches have simply repurposed the original source code to consider more than one chain. At the time the present work was begun, however, the only one of these methods to be available in final, published form was the Skolnick group’s method AF2Complex (Gao et al., 2022a) and it is this method, therefore, that has been used throughout the present work. AF2Complex provides several metrics for assessing the confidence of predicted protein complexes in addition to the original metrics provided by the underlying AlphaFold2 source code (Jumper et al., 2021), and it has already been applied to predict several new interactions for proteins that make up the cell wall of *E. coli* (Gao et al., 2022b).

Here, we use AF2Complex in combination with a number of other computational methods to build atomic models of the major TECs involved in lipid transport in *E. coli*. A comprehensive review of the TECs considered here has been reported recently (Giacometti et al., 2022). For some of these TECs, most of the “hard work” has already been performed by structural biologists using conventional experimental techniques, and the role played here by the computational methods is to fill in the remaining missing details. For others, there are significant gaps in our knowledge, and AF2Complex plays a potentially crucial role in identifying both the likely oligomeric states of some of the key proteins and in predicting the details of their interfaces with other proteins.

In applying AF2Complex to TECs, there are two major problems that must be addressed. The first stems from the sheer size of the complexes: three out of the four TECs considered here are sufficiently large in terms of their overall residue counts that an immediate application of AF2Complex is effectively precluded by RAM and/or computer time limitations. We attempt to circumvent this problem here by using a divide-and-conquer approach, making predictions for subcomplexes and then superimposing them to assemble complete models. The second major problem to overcome is the requirement that, in a TEC, both ends of the complex must be placed appropriately within membranes that are separated by ∼250 Å. Excellent computational methods are already available for assessing how a protein structure might be most favorably oriented within a membrane (Tusnády et al., 2004; Lomize et al., 2006; Lomize et al., 2022). Here we use the very popular OPM method (Lomize et al., 2022), applying it separately to the two ends of a TEC to predict how they are likely to be oriented within the IM and OM. But as we show here, the resulting predictions almost always result in orientations of the IM and OM that are not parallel with each other and that make it difficult to imagine, therefore, how the TECs might be properly placed in their cellular homes. To solve this issue, we have written code that, when combined with the popular molecular dynamics (MD) engine, GROMACS (Abraham et al., 2015), performs a series of restrained energy minimizations that progressively “morph” the TEC structures until their IM and OM regions are brought into alignment. We think that the combination of the previously reported AlphaFold2, AF2Complex, and OPM methodologies, together with the new “membrane morphing” protocol reported here, is likely to be a powerful method for building complete atomic models of a wide range of large protein complexes in the future.

## Results

In what follows, we describe the strategies that we have used to build complete atomic models of four key TECs involved in lipid transport in *E. coli*. For each complex, we begin by outlining what is already known about the complex, highlighting those parts or components that have been resolved by the experimental structural biology community. When needed, we next describe the use of AF2Complex in attempts to determine the oligomeric states of those components of the TECs for which structural data are absent. We then describe the use of AF2Complex to predict the structures of key subcomplexes within each TEC. In all cases, we annotate subcomplexes in the form A*n^i-j^*:B*m^k-l^*, where A and B are the gene names of the different proteins, *n* & *m* refer to their respective stoichiometries in the subcomplex, and *i-j* & *k-l* identify the range of residues from the full-length sequences that are included in the subcomplex prediction. We outline how the predicted structures of subcomplexes are combined to produce an initial complete model of the TEC, before describing how membrane morphing simulations are used to bring their IM and OM components into alignment in a final model.

Before proceeding, it is worth noting two general features of our use of AF2Complex. First, while several metrics have been proposed for estimating the reliability of structures predicted by AF2Complex, we focus here on ranking predictions by the <u>p</u>redicted interface <u>TM</u> (piTM) score proposed by the Skolnick group (Gao et al., 2022a; Gao et al., 2022b) because we found this to be generally the most reliable in the current setting. However, we also found that even this score can occasionally provide a misleading view of the plausibility of a predicted structure: in some cases, we obtained very favorable piTM scores from predicted structures that placed copies of the same chain directly on top of each other, i.e. in positions that would be impossible due to steric clashes. This issue has been noticed by others when using AlphaFold2-based methods (Gao et al., 2022a; David et al., 2022); the workaround to this problem that we adopted here was to only accept a prediction after it had been visually inspected: predictions containing truly egregious steric clashes due to erroneous chain superpositions are easy to identify and can easily be removed from further consideration. All of the piTM scores reported here, therefore, are for predicted structures that are free of non-physical chain superpositions.

The second key feature of our use of AF2Complex is that, for a number of the subcomplexes considered here, we repeated the predictions a number of times using different values assigned to the random seed. Repeating AlphaFold2 predictions for single-chain proteins is usually unnecessary, and implementations of AlphaFold2-based predictions typically restrict themselves to providing a single structure predicted by each of the five different neural network models optimized by the DeepMind team (i.e. five structures are returned from a single prediction submission). For some of the higher-order subcomplexes considered here, however, we found that the final quality of the predicted complexes often showed significant variability from run to run, with some of the neural network models producing greater variability than others. And, for one particular subcomplex prediction, we were fortunate to “strike gold” with a plausible model in only one out of 50 predicted structures.

### The Lpt TEC

The first TEC that we examine is the lipo<u>p</u>olysaccharide transport (Lpt) complex that transports molecules of lipopolysaccharide (LPS) from the IM to the OM (for a recent review, see Giacometti et al., 2022). Individual components of this complex have been solved independently by a number of groups using conventional experimental techniques (Suits et al., 2008; Qiao et al., 2014; Li et al., 2019; Figure 1A), and plausible images of the entire complex have already appeared in prior publications (e.g. Robinson, 2019; Giacometti et al., 2022), albeit with little information provided on how the structures shown in the images were constructed. The complete Lpt complex is known to contain: (a) an ABC transporter resident in the IM (subunits LptB_2_FGC), (b) a component resident in the OM (subunits LptDE), and (c) a “bridge” of LptA monomers that spans the periplasmic space, connecting LptC in the IM to LptD in the OM (Figure 1A). The presence of a LptA bridge is inferred in part from the observation that Lpts A, C and D all contain β jelly roll-like domains with exposed hydrophobic surfaces that have been shown to be important for LPS transport (Okuda et al., 2016). The number of LptA monomers required to bridge the periplasmic space has not yet been definitively established experimentally, but may be as high as five: importantly, LptA has been shown to form homo-oligomers (Suits et al., 2008).

**Figure 1.**
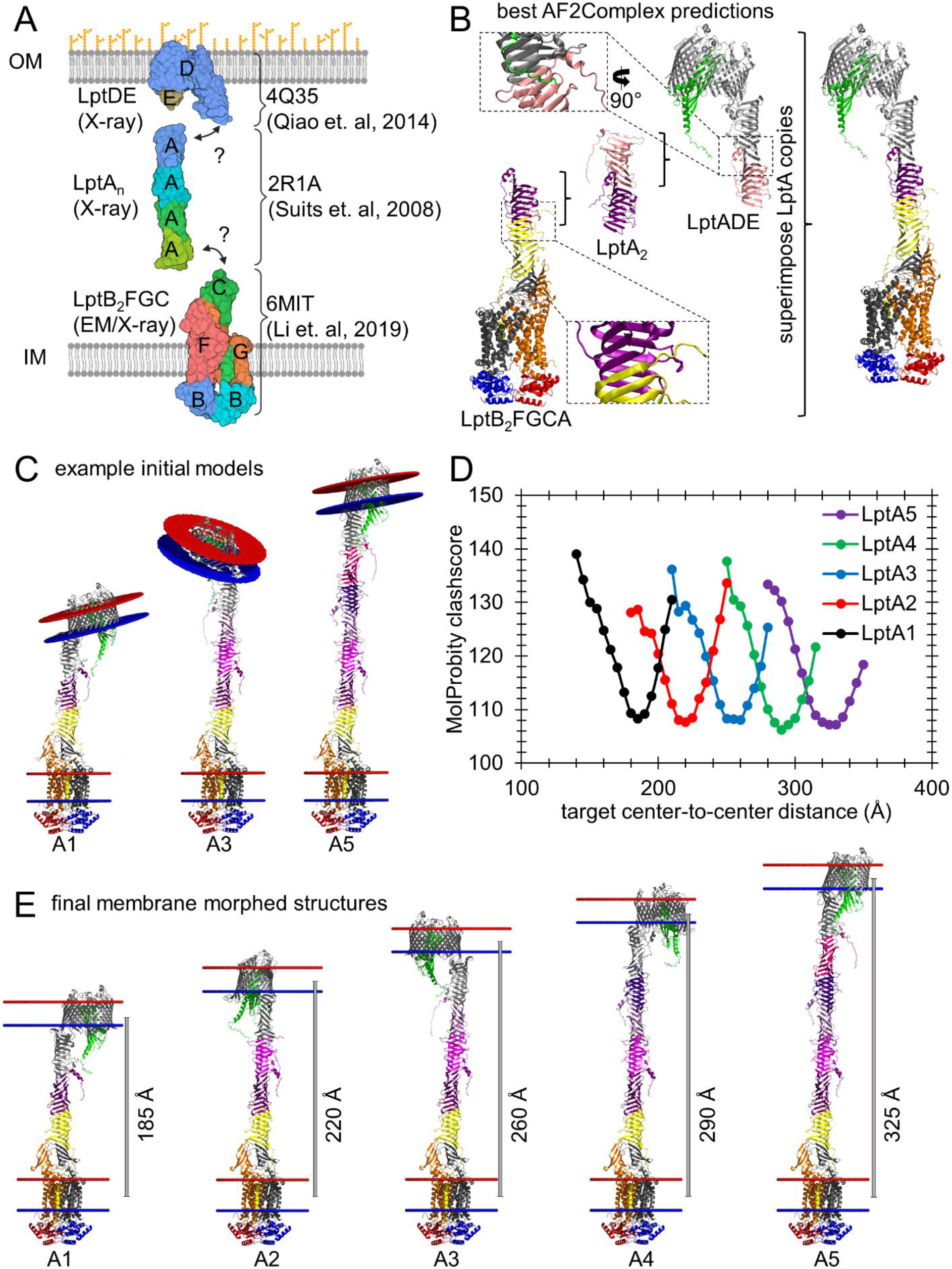
Building a complete atomic model of the Lpt TEC. (A) Schematic diagram of the Lpt TEC using solved structures labeled by RCSB IDs and their corresponding experimental method (created using BioRender). (B) Example AF2Complex predictions that were used to build complete Lpt complex models through superimpositions of LptA. Zoomed inserts highlight the LptA:LptC and LptA:LptD interfaces. (C) Examples of complete Lpt complex models built using superimpositions with increasing copies of LptA (colored various shades of purple) from left to right showing one (A1), three (A3) and five (A5) chains. (D) Plot of MolProbity clashscores versus the target center-to-center distance used as input in membrane morphing simulations (see main text) are shown for Lpt TEC models built with one (LptA1) to five (LptA5) copies of LptA. (E) Images of complete Lpt complex models with the lowest MolProbity clashscore after membrane morphing simulations. All inner membrane (IM) and outer membrane (OM) positions were predicted using OPM and are represented as circular planes of blue and red beads to show the inner and outer leaflet boundaries, respectively. All cartoon structure representations were generated in PyMol or VMD.

Depending on the number of LptA monomers that are included, the total number of residues in the Lpt complex ranges from 2,326 residues (for one LptA) to 2,958 (for five LptAs). Since these residue-counts are at or beyond the limit of what is generally feasible for AlphaFold2-based structure predictions (Jumper et al., 2021), we used a divide-and-conquer approach to predict atomic structures of subcomplexes and then sought to assemble them into a model of the entire complex (Figure 1B; see STAR Methods). To this end, we performed independent AF2Complex predictions for the following constructs: (a) LptB_2_FGCA, (b) LptA_2_ and (c) LptADE. Predictions for each of these subcomplexes allow us: (a) to assign plausible coordinates to those residues missing from the already-solved structures and (b) to predict the necessary interfaces that enable LptA to form a bridge between the IM and OM components of the complex.

For the three Lpt subcomplexes considered here, the highest piTM scores obtained from a single AF2Complex submission were all close to or above the threshold of 0.5 proposed by the Skolnick group (Gao et al., 2022a) as an indicator of a reliable prediction (Table 1). As anticipated, we found that the predictions with the highest piTM scores (Figure 1B) also nicely reproduced the structures of components found in experimentally solved structures. More interestingly, the predicted structures of the LptB_2_FGCA (IM) subcomplex and the LptADE (OM) subcomplex provide new, plausible atomic views of the probable LptA:LptC and LptA:LptD interfaces, respectively. In both cases, the interfaces involve antiparallel alignment of β-strands from neighboring chains (Figure 1B zoomed inserts); this arrangement matches well with what was proposed previously in the literature (Suits et al., 2008; Sperandeo et al., 2011; Freinkman et al., 2012; Okuda et al., 2016).

**Table 1.** Summary statistics for all AF2Complex predictions. A summary of statistics for all AF2Complex predictions performed in this work are grouped by the TEC system and show the stoichiometry of all subunits included in the predictions, the total residue counts, the total number of prediction replicates, the piTM scores of the best predictions, the neural networks responsible for the best predictions, and the cellular localization of the predictions.

| TEC system | prediction subunit stoichiometry | total prediction residue count | total number of predictions | best prediction piTM score | best prediction neural network | predicted cellular localization |
| --- | --- | --- | --- | --- | --- | --- |
| Lpt | B<sub>2</sub>FGCA | 1557 | 5 | 0.48 | 3 | IM |
|  | A<sub>2</sub> | 316 | 5 | 0.69 | 3 | P |
|  | DEA | 1093 | 5 | 0.73 | 3 | OM |
| Pqi | A<sub>2</sub> | 834 | 50 | 0.05 | 3 | IM |
|  | A<sub>3</sub> | 1251 | 50 | 0.06 | 3 | IM |
|  | AB<sub>6</sub><sup>1-133</sup> | 1215 | 50 | 0.55 | 5 | IM |
|  | A<sub>2</sub>B<sub>6</sub><sup>1-133</sup> | 1632 | 50 | 0.42 | 5 | IM |
|  | A<sub>3</sub>B<sub>6</sub><sup>1-133</sup> | 2049 | 50 | 0.29 | 2 | IM |
|  | B<sub>6</sub><sup>41-412</sup> | 2232 | 5 | 0.44 | 3 | P |
|  | B<sub>6</sub><sup>285-546</sup> | 1572 | 5 | 0.49 | 2 | P |
|  | B<sub>6</sub><sup>432-546</sup>C<sub>6</sub> | 1812 | 50 | 0.56 | 2 | P/OM |
|  | C<sub>2</sub> | 374 | 5 | 0.68 | 5 | OM |
|  | C<sub>3</sub> | 561 | 5 | 0.78 | 5 | OM |
|  | C<sub>4</sub> | 748 | 5 | 0.79 | 5 | OM |
|  | C<sub>6</sub> | 1122 | 5 | 0.62 | 2 | OM |
|  | C<sub>8</sub> | 1496 | 5 | 0.51 | 3 | OM |
|  | C<sub>12</sub> | 2244 | 5 | 0.47 | 3 | OM |
| Let | A<sub>2</sub> | 854 | 50 | 0.25 | 2 | IM |
|  | A<sub>3</sub> | 1281 | 50 | 0.16 | 2 | IM |
|  | AB<sub>6</sub><sup>1-137</sup> | 1249 | 50 | 0.60 | 2 | IM/P |
|  | A<sub>2</sub>B<sub>6</sub><sup>1-137</sup> | 1676 | 50 | 0.41 | 3 | IM/P |
|  | A<sub>3</sub>B<sub>6</sub><sup>1-137</sup> | 2103 | 50 | 0.29 | 4 | IM/P |
| Tam | AB | 1836 | 50 | 0.48 | 3 | IM/P/OM |

IM = inner membrane
OM = outer membrane
P = periplasm

With plausible predicted atomic models for the three subcomplexes in hand, we constructed models of the complete Lpt complex by superimposing LptA chains shared between neighboring subcomplexes (see STAR Methods). Figure 1C shows examples of initial models built in this way that contain between one, three, and five LptA chains, respectively. In all cases, the entire LptC- LptA_1_-…-LptA_n_-LptD assembly can be viewed as comprising a single, continuous β-sheet structure that stretches all the way from the IM to the OM. Although the models shown in Figure 1C are complete, however, they are unlikely to be very good models for how the Lpt complex is likely to appear *in vivo*. The reason for this can be seen from inspecting the relative orientations of the IM and OM predicted for these models by the membrane orientation algorithm OPM (Figure 1C; Lomize et al., 2022). In all cases, the predicted orientations of the two membranes are tilted substantially with respect to each other, making it difficult to see how the TEC could be situated across the periplasm without significantly distorting one or both membranes.

To resolve this issue, we devised a molecular mechanics simulation-based procedure that seeks to minimally adjust the models to bring the planes of their predicted OM and IM regions into parallel alignment (see STAR Methods). Throughout this simulation protocol, which we term “membrane morphing”, atoms predicted by OPM to be embedded in either the IM or the OM are subjected to position restraints that act to maintain their depth within each membrane, but the plane of the OM is progressively adjusted until it becomes parallel to the plane of the IM. All other atoms in the TEC remain unrestrained during the procedure, and the entire complex is held together by a simple generic energy model that seeks to preserve bonds, angles, dihedral angles, and close inter-atomic contacts at their initial values (see STAR Methods). Since the membrane morphing procedure makes use of the popular GROMACS molecular simulation code (Abraham et al., 2015), it is fast and reliable, and only requires a single major input value from the user, which is the desired final separation distance between the centers of the IM and OM.

Since literature estimates of the periplasmic width vary significantly (Graham et al., 1991; Oliver 1996; Matias et al., 2003; Cohen et al., 2017), we conducted a number of membrane morphing simulations to build final Lpt models covering a wide range of possible center-to-center distances between the two membranes. To identify the most plausible center-to-center distance for each variant of the Lpt complex, we made use of the Richardson group’s MolProbity webserver (Davis et al., 2007; Chen et al., 2010; Williams et al., 2018) which assesses the quality of protein atomic structures (see STAR Methods). While the MolProbity server reports several quality measures, we chose to focus on the “clashscore” (a measure of the number of bad steric clashes) since this is expected to identify serious problems in the structures that may prevent their use in downstream studies. Importantly, for all variants of the Lpt complex, we found that the clashscore exhibited a roughly parabolic dependence on the center-to-center distance between the two membranes, with an optimal value being easy to identify in each case (Figure 1D). The optimal center-to-center distances identified from this analysis matched closely with our subjective assessments based on visual inspection of the final models. Figure 1E shows all of the final Lpt models obtained after membrane morphing and after a final short energy minimization in GROMACS; movies showing all five morphing trajectories can be found in Movies S1-S5. For the Lpt models containing 1, 2, 3, 4, and 5 LptA monomers the optimal center-to-center distances between the IM and OM were 185, 220, 260, 290 and 325 Å, respectively; the addition of each LptA monomer, therefore, causes an increase of ∼35 Å in the preferred distance. To our knowledge, these models provide the first complete atomic structures of the Lpt complex with LptA bridging the IM and OM components.

### The Pqi TEC

The next system that we consider is the <u>p</u>ara<u>q</u>uat-inducible (Pqi) system that has been implicated in trafficking lipids between the IM and OM (for a recent review, again see Giacometti et al., 2022). A schematic representation of the known components of this system is shown in Figure 2A. PqiB forms a hexameric channel that is thought to directly connect an IM-resident component, PqiA, with a periplasm-facing, OM-anchored lipoprotein, PqiC (Giacometti et al., 2022). CryoEM studies have revealed the molecular architecture of a substantial fraction of the PqiB channel: most of residues 42-431 of the 546-residue protein have been resolved, revealing that the hexameric channel is composed of three rings of stacked <u>m</u>ammalian <u>c</u>ell <u>e</u>ntry (MCE) domains followed by a narrow “needle” formed by entwined α-helices (Ekiert et al., 2017; Isom et al., 2020). The end-to-end length of the most complete cryoEM PqiB structure deposited in the RCSB by the Vale group (RCSB ID: 5UVN), is ∼150 Å, which is some distance short of estimates of the periplasmic width (Graham et al., 1991; Oliver et al., 1996; Vollmer & Seligman 2010). It seems reasonable to assume, therefore, that the 115 C-terminal residues of PqiB that are not resolved in the cryoEM structure might serve to bridge the gap. To this end, the Vale group proposed a compelling model in which the needle is extended by a further ∼70 Å through the addition of intertwined poly-alanine helices (Ekiert et al., 2017); the latter model is shown in Figure 2B.

**Figure 2.**
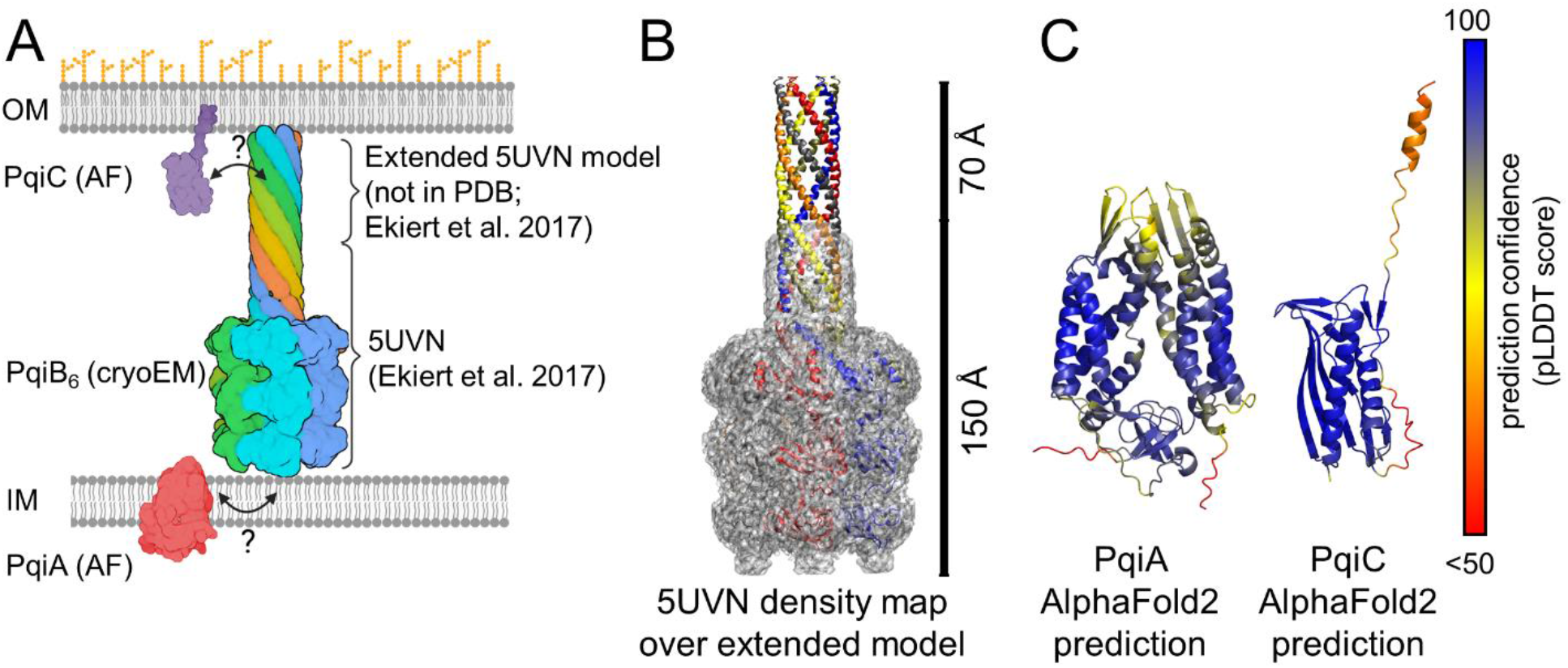
Known components of the Pqi TEC. (A) A schematic diagram of the Pqi TEC is shown using both solved components that are labeled by either RCSB IDs with the corresponding experimental method or as AlphaFold (AF) models for unsolved components (created using BioRender). (B) The PqiB_6_ cryo-electron microscopy density map and a corresponding extended model (RCSB ID: 5UVN; Ekiert et al. 2017) are shown as a grey surface and colored cartoon chains, respectively. (C) Cartoon representations of the AlphaFold2 predictions for PqiA and PqiC are shown colored by prediction confidence per residue (pLDDT score) on a spectrum of red – yellow – blue for unconfident – moderate confidence – high confidence scores, respectively. All structure representations were generated in either PyMol or VMD.

With regard to the other members of the complex, according to the SEQATOMS Blast server (Brandt et al., 2008) PqiA has no high sequence-identity match with any structure in the RCSB; PqiC, however, matches with a sequence identity of 84% to an uncharacterized protein from *Enterobacter cloacae* (RCSB ID: 6OSX; Chang et al., 2019 unpublished). Although there is no strong match in the RCSB for PqiA, the AlphaFold Protein Structure Database hosted by the EMBL-EBI (https://alphafold.ebi.ac.uk; Varadi et al., 2022) contains high-confidence predictions for both the monomeric forms of both PqiA and PqiC (Figure 2C). This suggests that completing a model of the entire Pqi complex using AF2Complex predictions is likely to be feasible.

Construction of a model of the complete Pqi TEC was carried out in the following stages. First, we used AF2Complex to attempt to determine the probable oligomeric states of the IM-bound PqiA, and the OM-bound PqiC. Second, we used AF2Complex to predict structures for the PqiA:PqiB and PqiB:PqiC interfaces, with fragments of PqiB being used in both predictions rather than the full-length protein owing to the high residue counts involved. Third, we used structural superpositions to piece together four overlapping fragments of PqiB into an initial model of the complete Pqi TEC. Finally, we used membrane morphing simulations to bring the predicted planes of the IM and OM into alignment with each other to produce a final model.

Our first goal was to resolve the likely oligomeric state of PqiA. One crude way that can be used to gauge the range of possible oligomeric states is to examine the copy numbers of the Pqi proteins estimated by the Weissman group from their ground-breaking ribosome profiling experiments on *E. coli* (Li et al., 2014; Table S1). For growth in rich-defined media, ribosome profiling leads to estimated copy numbers of 143, 439, and 612 per cell for PqiA, PqiB, and PqiC respectively; for growth in 0.2% glucose media, the estimated copy numbers are 34, 96, and 175, respectively. Since PqiB is known experimentally to be a hexamer (see above), it seems reasonable to imagine from these numbers that PqiA could be a monomer, a dimer, or a trimer, but is unlikely to be anything much larger. The AlphaFold2-predicted structure of the PqiA monomer (Figure 2C) has a confident average pLDDT score of 87.6 ± 10.6, and importantly, it also has a favorable predicted membrane transfer free energy of -67.2 kcal/mol as computed by OPM, suggesting that the monomer alone is a viable model of an IM-resident protein. However, to see if there was any evidence of a subunit-subunit interface that might drive homo-oligomerization, we used AF2Complex to predict the structures of possible dimeric and trimeric forms (i.e. PqiA_2_ and PqiA_3_, respectively). For both oligomeric forms, the best piTM scores obtained were less than 0.1 (see Figure 3A and Table 1) with very few atomic contacts occurring between chains in these structures (see Figure S1 for the predicted structures with the highest piTM scores for PqiA_2_ and PqiA_3_). Taken together, these results provide no compelling evidence for the existence of a stable interaction between subunits, and so we tentatively concluded that PqiA is likely to be a monomer.

**Figure 3.**
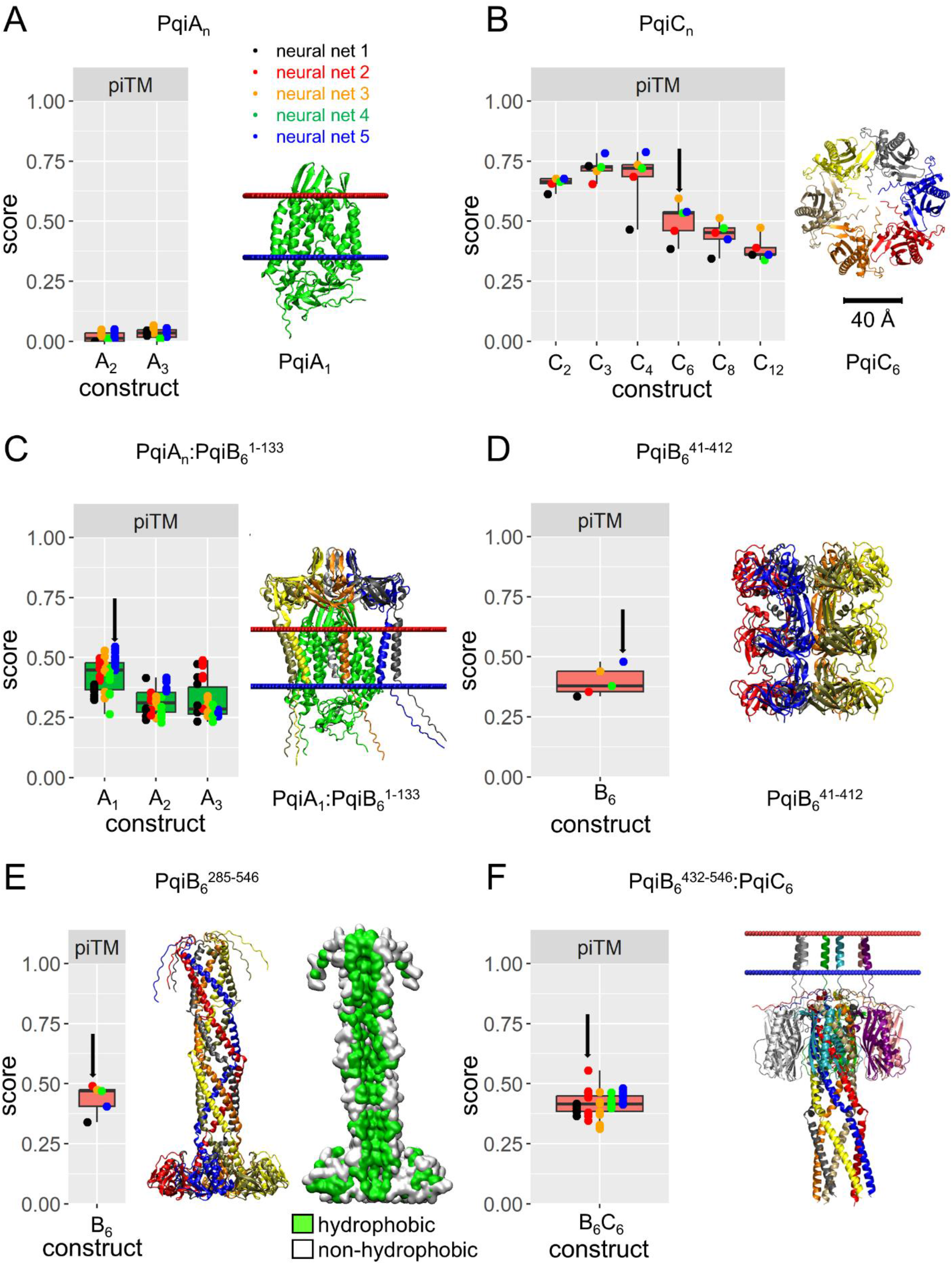
AF2Complex predictions of Pqi TEC components. Each section shows the results from AF2Complex predictions performed on Pqi TEC components through a box plot of the AF2Complex piTM scores and the structure of the best prediction (indicated by a black arrow in the box plot where applicable). (A) Plot of the piTM scores for all PqiA_2_ and PqiA_3_ predictions and the PqiA AlphaFold2 prediction placed in a Gram-negative bacterial IM using OPM. (B) Plot of the piTM scores for all PqiC homo-oligomeric predictions and the PqiC_6_ prediction with the highest piTM score. (C) Plot of the piTM scores for all PqiA_n_:PqiB_6_^1-133^ predictions and the PqiA_1_:PqiB_6_^1-133^ prediction with the highest piTM score placed in a Gram-negative bacterial IM using OPM. (D) Plot of the piTM scores for all PqiB_6_^41-412^ predictions and the prediction with the highest piTM score. (E) Plot of the piTM scores for all PqiB_6_^285-546^ predictions and two representations of the prediction with the highest piTM score: one using a cartoon representation and another using a surface representation showing the hydrophobic (green) and non-hydrophobic (white) residues. (F) Plot of the piTM scores for all PqiB_6_^432-546^:PqiC_6_ predictions and the prediction with the highest piTM score placed in a Gram-negative bacterial OM using OPM. All box plots show individual piTM scores as points colored by the neural network responsible for the prediction and were generated in R (see Methods), while all structure representations were generated in either PyMol or VMD and show the structure colored by chain with the inner and outer leaflet of the membrane shown as blue and red spheres, respectively.

We followed a similar strategy to identify a plausible oligomeric state for PqiC. The AlphaFold2- predicted structure of monomeric PqiC (Figure 2C) has an average pLDDT score of 89.3 ± 14.4. As noted above, the SEQATOMS Blast server indicates that PqiC has a very strong match to RCSB ID 6OSX; this structure is assigned as a monomer both by the authors who deposited it and by the PQS webserver, which attempts to identify oligomeric states on the basis of crystal contacts (Henrick & Thornton 1998). Interestingly, however, the Weissman group’s relative copy numbers for PqiB and PqiC (see above) strongly suggest that PqiC is likely to be oligomeric. Since PqiC’s smaller size makes it possible for us to consider a wider range of potential oligomeric states, we used AF2Complex to predict structures for PqiC_2_, PqiC_3_, PqiC_4_, PqiC_6_, PqiC_8_, and PqiC_12_. In contrast to what we observed with PqiA, there were strong indications of favorable interactions between subunits in all of the predicted oligomeric states. When considered in order of increasing oligomer size, the piTM scores of these predictions initially increase, reaching a maximum value with PqiC_4_, before decreasing as the oligomers became larger (see Figure 3B and Table 1). However, none of the predicted structures for PqiC_2_, PqiC_3_, and PqiC_4_ form a closed ring structure that would be compatible with the known ring structure of PqiB_6_ (see Figure S2 for the predicted structures with the highest piTM scores of all PqiC oligomers). We therefore chose to reject these predictions and selected instead PqiC_6_ as the most likely oligomeric state since, of the oligomers that successfully formed closed rings, it produced the highest piTM score. Interestingly, this choice of oligomeric state not only matches that of PqiB_6_, it is also further supported by the fact that the predicted PqiC_6_ ring structure has an internal diameter (∼40 Å) that matches well with the external diameter of the PqiB_6_ needle (∼35 Å from RCSB ID: 5UVN; Ekiert et al., 2017).

We next set about identifying a series of four subcomplexes whose structures could be predicted with AF2Complex, and which could then be superimposed upon each other to assemble the complete Pqi TEC. Each of these four subcomplexes involved some kind of fragment of PqiB_6_. The first was intended to model the PqiA:PqiB interface. Our earlier predictions of PqiA alone had suggested that it is likely to be a monomer, but to leave other options open we also considered the possibility that PqiA might become dimeric or trimeric when interacting with PqiB_6_. The predicted piTM scores obtained for the monomeric PqiA_1_:PqiB_6_^1-133^ construct were consistently higher than those of the PqiA_2_:PqiB_6_^1-133^ and PqiA_3_:PqiB_6_^1-133^ constructs (see Figure 3C and Table 1), further supporting the idea that PqiA is likely to be monomeric (see Figure S3 for the predicted structures with the highest piTM scores of PqiA_2_:PqiB_6_^1-133^ and PqiA_3_:PqiB_6_^1-133^). Notably, in the best-scoring models of PqiA_1_:PqiB_6_^1-133^, the PqiA monomer is surrounded by N-terminal transmembrane helices contributed by each of the six PqiB chains (Figure 3C). The combined membrane transfer free energy of the PqiA_1_:PqiB_6_^1-133^ construct is predicted to be -177.0 kcal/mol by OPM, which again indicates that it is a plausible model for an IM-resident subcomplex.

The next two subcomplexes predicted were intended to complete missing details of the PqiB hexamer. Although most of the PqiB hexamer was solved in the cryoEM 5UVN structure (Figure 2B; Ekiert et al., 2017), coordinates for residues 348-354 and 388-401 are absent, and all residues from 432-546 are unresolved. We performed two AF2Complex predictions on hexameric constructs covering PqiB residue ranges from 41-412 and 285-546 in order to fill in these missing residues. The results for the PqiB_6_^41-412^ construct (Figure 3D) are comparatively uninteresting in the sense that they simply recapitulate what is already known from the cryoEM structure. Much more interesting and impressive, however, are the results for the PqiB_6_^285-546^ construct (Figure 3E). In these predictions, AF2Complex extends the helical needle observed originally in 5UVN with a further ∼90 residues (Figure 3E, PqiB_6_^285-546^) and in a manner that bears a striking similarity to the extended poly-alanine helices modeled in by the Vale group (Figure 2B). Moreover, the physicochemical characteristics of the predicted needle interior are consistent with the Pqi TEC’s role in transferring lipids: the interior surface of the needle extension is formed almost exclusively by hydrophobic sidechains that protrude into it (Figure 3E).

The fourth and final subcomplex that we predicted was focused on modeling the PqiB:PqiC interface. For these predictions we used the construct PqiB_6_^432-546^:PqiC_6_ with the PqiB construct being selected as it contains all the residues necessary to form the top end of the completed needle seen in our earlier predictions of PqiB alone (see above; PqiB_6_^285-546^). Predictions for this subcomplex were unusual in that the very best model had a piTM score that was notably higher than the other scores (Figure 3F arrow). Importantly, in that predicted structure the PqiB hexameric needle was neatly threaded through the center of the hexameric PqiC ring, with the N-terminal residues of each PqiC chain, which in reality serve as a “lipobox” signal peptide (Nakayama & Zhang-Akiyama, 2017), strikingly oriented toward the expected position of the OM (Figure 3F). Evidently, AlphaFold2 appears to somehow “know” that signal peptides, even when they are destined to be cleaved and replaced with a tripalmitoylated cysteine, should be oriented normal to the membrane. Predictions with lower piTM scores typically contained complete hexameric PqiB needles and/or hexameric PqiC rings but did not assemble them into a single symmetric complex (see Figure S4 for examples of some of these “misfires”). It appears, therefore, that correct assembly of this particular subcomplex with AF2Complex requires a coordination of events during the calculations that occurs only infrequently.

Finally, we built a complete model of PqiA_1_:PqiB_6_:PqiC_6_ by superimposing residues shared in each subcomplex prediction (Figures 4A and 4B). We then used membrane morphing simulations to bring the predicted orientations of the IM and OM into alignment and found that a center-to-center IM-OM distance of 265 Å resulted in the best MolProbity clashscore (Figure 4C). The final structure obtained after a short energy minimization in GROMACS is shown in Figure 4D, and a movie showing the membrane morphing is provided in Movie S6.

**Figure 4.**
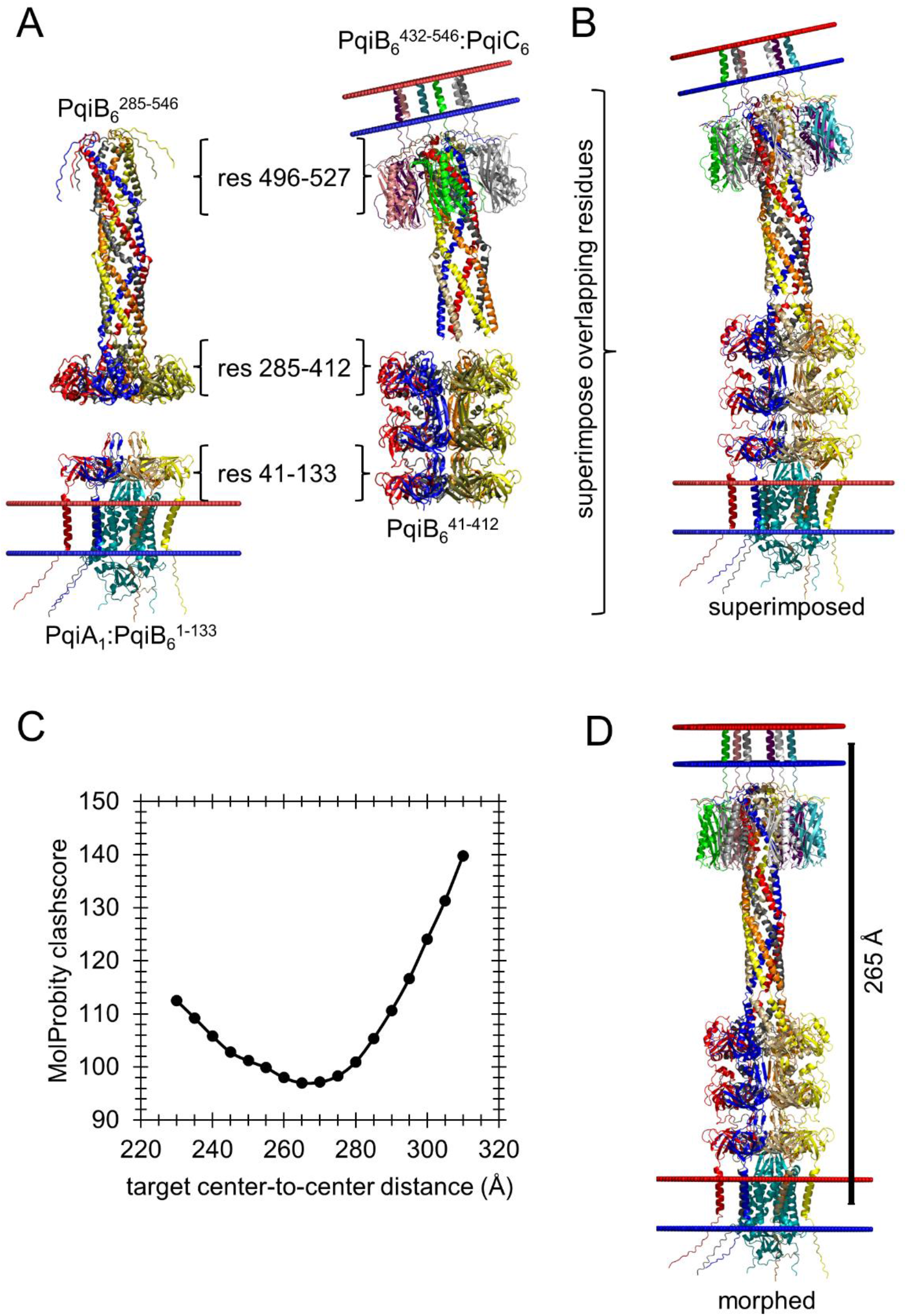
Modeling the complete Pqi TEC . (A) AF2Complex predictions with the highest piTM scores are shown for the constructs PqiA_1_:PqiB_6_^1-133^, PqiB_6_^41-412^, PqiB_6_^285-546^ and PqiB_6_^432- 546^:PqiC_6_. The residues used to superimpose each construct are shown within brackets that point to their relative locations in the structures. (B) A complete model of the Pqi TEC built using superimpositions (see Methods). (C) Plot of MolProbity clashscores versus the target center-to-center distances used as input in membrane morphing simulations of the complete Pqi TEC. (D) The final structure of the Pqi TEC membrane morphing simulation that has the lowest MolProbity clashscore. All cartoon structural representations were generated in either VMD or PyMol and show the structure colored by chain with the inner and outer leaflet of the membrane shown as blue and red spheres, respectively.

### The Let TEC

The next TEC we consider is the lipophilic <u>e</u>nvelope-spanning tunnel (Let) system, previously known as the Yeb system (Isom et al., 2020). The Let complex is composed of an IM-resident component, LetA, and an IM-anchored periplasm-spanning protein, LetB, which together act to maintain membrane integrity alongside the Pqi system (Nakayama & Zhang-Akiyama, 2017). There has been some discussion in the literature whether there might be a third component of the complex resident in the OM (Nakayama & Zhang-Akiyama, 2017) but at the time of writing no candidate appears to have been definitively identified. Figure 5A shows what is currently known regarding the Let system: the structure and oligomeric state of LetA remain unresolved, but the structure of LetB has been well characterized experimentally. In particular, recent cryoEM studies have revealed that LetB is anchored to the IM through a single-pass transmembrane helix at its N-terminus (Isom et al., 2017) which is followed by seven stacked MCE domains that extend ∼210 Å, forming a hexameric channel that surrounds a central hydrophobic tunnel (Ekiert et al., 2017; Liu et al., 2020; Isom et al., 2020; Vieni et al., 2022). The most complete of the structures resulting from these studies is shown in Figure 5B (RCSB ID: 6V0C; Isom et al., 2020).

**Figure 5.**
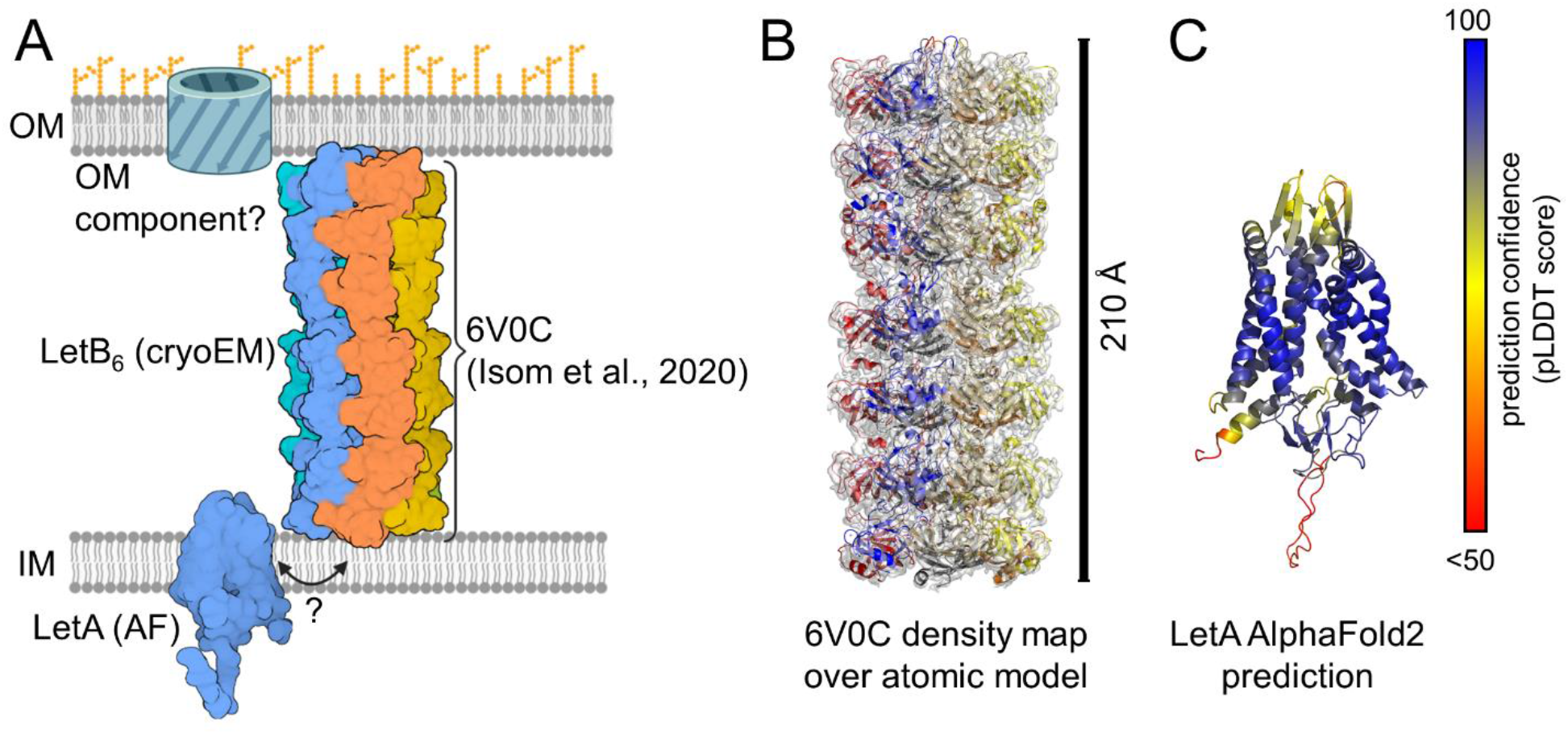
Known components of the Let TEC. (A) A schematic diagram of the Let TEC is shown using both solved components that are labeled by either RCSB IDs with the corresponding experimental method or as AlphaFold (AF) models for unsolved components (created using BioRender). (B) The cryo-electron microscopy density map and the corresponding atomic model for the LetB hexamer (RCSB ID: 6V0C) are shown as a grey surface and colored cartoon chains, respectively. (C) The AlphaFold2 prediction for LetA colored by prediction confidence per residue (pLDDT score) on a spectrum of red – yellow – blue for unconfident – moderate confidence – high confidence scores, respectively. All structure representations were generated in either PyMol or VMD.

Our first goal was to resolve the likely oligomeric state of LetA. Again, we began by looking at the protein copy numbers reported by the Weissman group (Li et al., 2014; Table S1). For growth in rich-defined media, the estimated copy numbers are 47 and 126 per cell for LetA and LetB, respectively; for growth in 0.2% glucose media, the estimated copy numbers are 35 and 77, respectively. Since LetB is known experimentally to be a hexamer (see above), it again seems reasonable to imagine from these numbers that LetA could conceivably be a monomer, a dimer, or a trimer, but is unlikely to be anything much larger. Although the structure of LetA has yet to be solved and there is no high sequence-identity match in the RCSB, the AlphaFold2 database again contains a high-confidence structural model with an average pLDDT score of 86.3 ± 12.2 (Figure 5C). Interestingly, this structure bears a strong similarity to AlphaFold2’s predicted structure of PqiA (Figure 2C) which is consistent with the fact that the sequence identity of the two proteins is 29%. As was the case with PqiA, the AlphaFold2-predicted monomer structure has a favorable predicted membrane transfer free energy of -53.6 kcal/mol as computed by OPM; this suggests again that the monomer is a viable model of an IM-resident protein.

To see if there was any evidence of a subunit-subunit interface that might drive homo-oligomerization, we used AF2Complex to predict the structures of possible dimeric and trimeric forms (i.e. LetA_2_ and LetA_3_, respectively). For both oligomeric forms, the best piTM scores obtained were 0.25 or lower (see Figure 6A and Table 1); the modest best scores obtained for these interactions appear to result from a β-sheet interaction that seems implausible for an IM-resident component (see Figure S5). As was the case with PqiA, therefore, these results provide no compelling evidence for the existence of a stable interaction between subunits, and so we tentatively concluded that LetA is likely to be a monomer.

**Figure 6.**
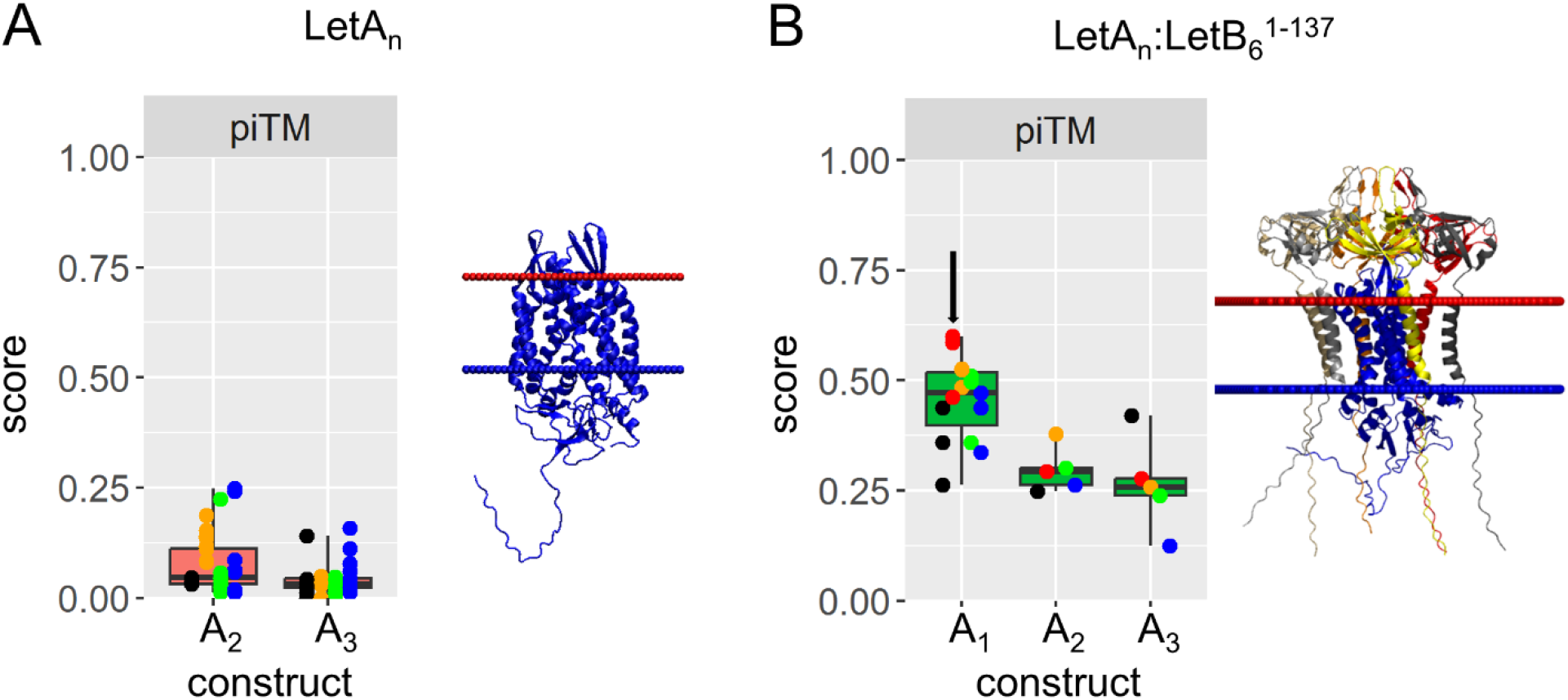
Modeling the LetA:LetB interface. (A) AF2Complex results for LetA homo-oligomeric structure predictions. On the left is a box plot of the AF2Complex piTM scores for LetA_2_ and LetA_3_ predictions and on the right is the LetA AlphaFold2 structure inserted into a Gram-negative bacterial IM using OPM. (B) AF2Complex results for LetA_n_:LetB_6_^1-137^ predictions. On the left is a box plot of the AF2Complex piTM scores for LetA_1_:LetB_6_^1-137^ (“A1”), LetA2:LetB_6_^1-137^ (“A2”), and LetA3:LetB_6_^1-137^ (“A3”) and on the right is the LetA_1_:LetB_6_^1-137^ prediction with the highest piTM score (indicated by the black arrow in the plot). All box plots show individual piTM scores as points colored by the neural network responsible for the prediction and were generated in R (see Methods), while all structure representations were generated in either PyMol or VMD and show the structure colored by chain with the inner and outer leaflet of the membrane shown as blue and red spheres, respectively.

Our next goal was to model the LetA:LetB interface. While our earlier predictions of LetA alone had suggested that it is likely to be a monomer, to leave other options open we again considered the possibility that it might become dimeric or trimeric when interacting with LetB_6_. The predicted piTM scores obtained for the monomeric LetA_1_:LeB_6_^1-137^ construct were consistently higher than those of the LetA_2_:LeB_6_^1-137^ and LetA_3_:LeB_6_^1-137^ constructs, further supporting the idea that LetA is likely to be monomeric (Figure 6B and Figure S6). Again, in the best-scoring models in which LetA was modeled as a monomer it is surrounded by N-terminal transmembrane helices contributed by each of the six LetB chains (Figure 6B). The combined membrane transfer free energy of the LetA_1_:LetB_6_^1-137^ construct is predicted to be -136.2 kcal/mol by OPM, which again indicates that it is a plausible model for an IM-resident subcomplex.

Finally, we built a complete model of the Let TEC by superimposing our best LetA_1_:LetB_6_^1-137^ prediction onto the LetB_6_ cryoEM structure (Figure 7A). The initial model obtained in this way has plausible interfaces but the predicted planes of the IM and OM are significantly tilted relative to each other (Figure 7B). We then used membrane morphing simulations to bring the predicted orientations of the IM and OM into alignment and found that a center-to-center IM-OM distance of 235 Å resulted in the best MolProbity clashscore (Figure 7C). The final model obtained after a short energy minimization in GROMACS is shown in Figure 7D; a movie illustrating the membrane morphing trajectory is provided in Movie S7.

**Figure 7.**
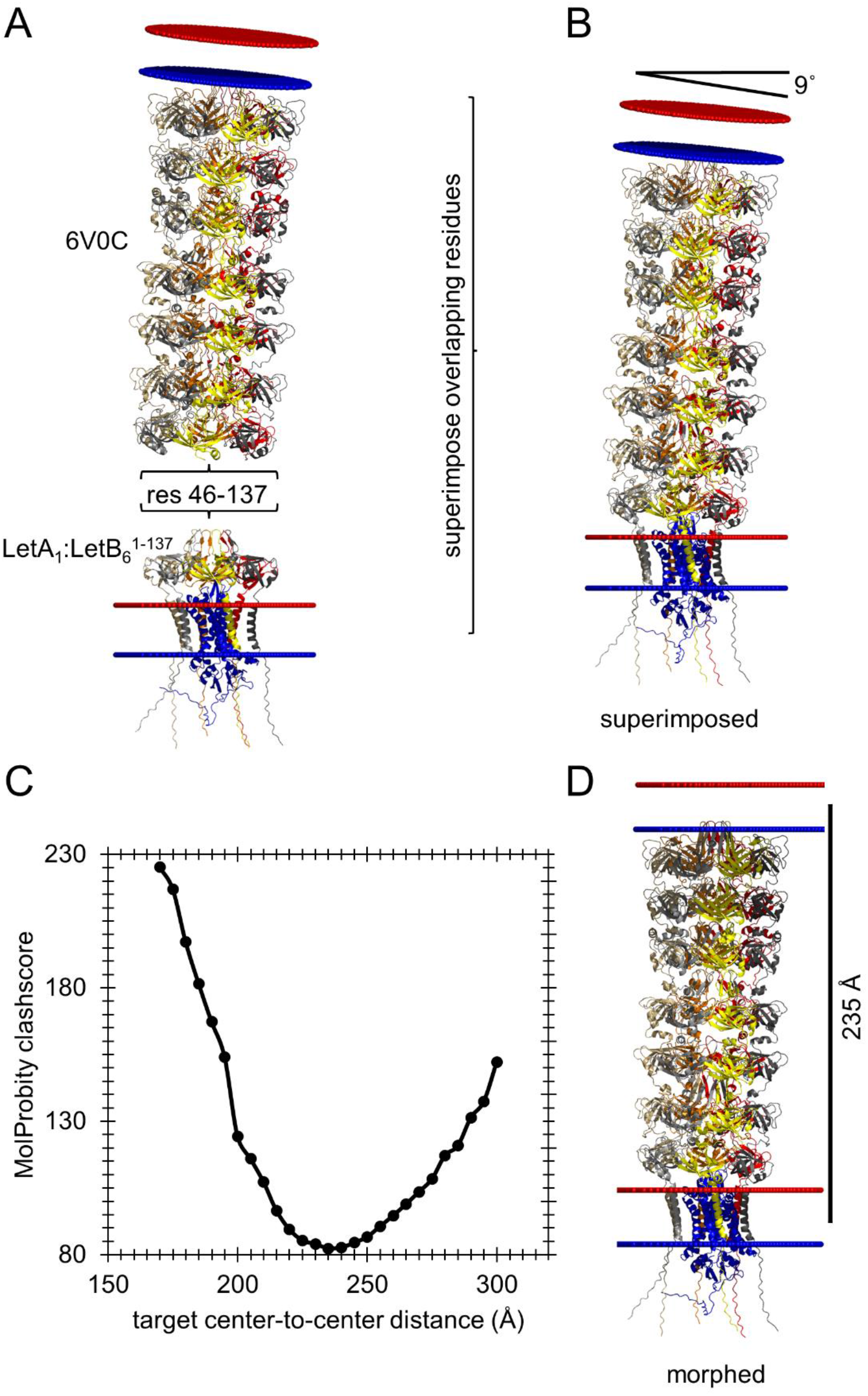
Modeling the complete Let TEC. (A) The LetA_1_:LetB_6_^1-137^ AF2Complex prediction with the highest piTM score (bottom) and the solved LetB_6_ structure (top, RCSB ID: 6V0C) are shown with the residues used in superimpositions to build a complete model indicated within brackets. (B) The complete Let TEC model built from superimposing residues 46-137 in both structures from (A). (C) Plot of MolProbity clashscores versus the target center-to-center distances used as input in membrane morphing simulations of the complete Let TEC. (D) Final structure of the Let TEC membrane morphing simulation that had the lowest MolProbity clashscore (target center-to-center distance of 235 Å). All structure representations were generated in either PyMol or VMD and show the structure colored by chain with the inner and outer leaflet of the membrane shown as blue and red spheres, respectively.

### The Tam TEC

While the above Lpt, Pqi, and Let systems are the most commonly cited lipid transport trans-envelope complex systems in *E. coli*, there are others that have also been shown to help maintain OM integrity and are thought to have lipid transport functionality. Chief among these systems are the six AsmA-like proteins (AsmA, TamB, YdbH, YicH, YhjG and YhdP). Very recently, AlphaFold2 predictions coupled with molecular biology experiments have suggested that these proteins transport lipids to the OM through extended hydrophobic core regions similar to that found in the Lpt system (Ruiz et al., 2021; Douglass et al., 2022). Mutational studies and comparisons with homologous protein systems suggest that five of these proteins are likely to be monomeric (Ruiz et al., 2021; Douglass et al., 2022), so that the existing AlphaFold2 predictions are likely sufficient to model these systems. The sixth member of the family, TamB, however, is known to be functionally dependent on forming a complex with the β-barrel OM protein, TamA (Selkrig 2012; Ruiz et al., 2021).

Figure 8A shows what is currently known about the Tam TEC. The structure of TamA has already been solved experimentally (RCSB ID: 4C00; Gruss et al., 2013; missing residues 19-24, 84-92). In contrast, only residues 963-1138 of TamB have been solved experimentally (RCSB ID: 5VTG; Josts et al., 2017), and no other region of the protein produces a significant match using the SEQATOMs BLAST server. The AlphaFold2 model for TamA (Figure 8B) has an average pLDDT score of 92.9 ± 12.0; the corresponding model for TamB (Figure 8C) also has an average pLDDT score of 87.9 ± 12.4. Together, these results suggest that a prediction of the complete Tam TEC is worth pursuing. Experimental characterization of the Tam TEC (Selkrig et al., 2012) has demonstrated it to have a 1:1 stoichiometry and the resulting residue count is sufficiently low that it is feasible to predict the structure of the entire complex using AF2Complex. We found that the prediction with the highest piTM score (0.48; Table 1) produced a plausible structure of the complex (Figure 9A), in which the interface between the two proteins forms through an interaction in which TamB serves to complete the β-barrel of TamA (Figure 9A zoomed insert). Importantly, this mode of interaction is in line with a previous suggestion about how these proteins might interact (Selkrig et al., 2012), indicating that the model predicted by AF2Complex is a plausible initial model of the Tam TEC.

**Figure 8.**
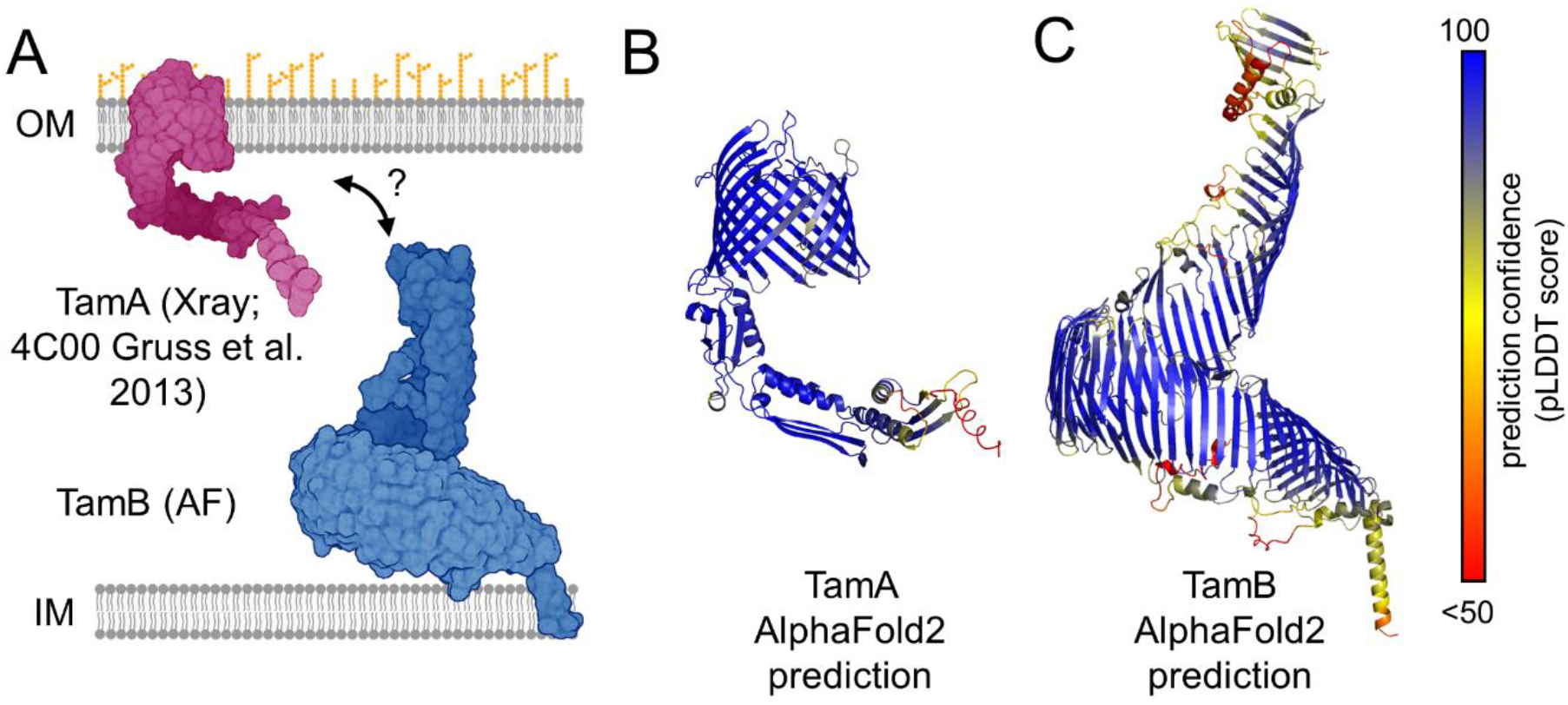
Known components of the Tam TEC. (A) A schematic diagram of the Tam TEC is shown using both solved components that are labeled by either RCSB IDs with the corresponding experimental method or as AlphaFold (AF) models for unsolved components (created using BioRender). AlphaFold2 predictions for (B) TamA and (C) TamB are represented as cartoons and are colored by prediction confidence per residue (pLDDT score) on a spectrum of red – yellow – blue for unconfident – moderate confidence – high confidence scores, respectively.

**Figure 9.**
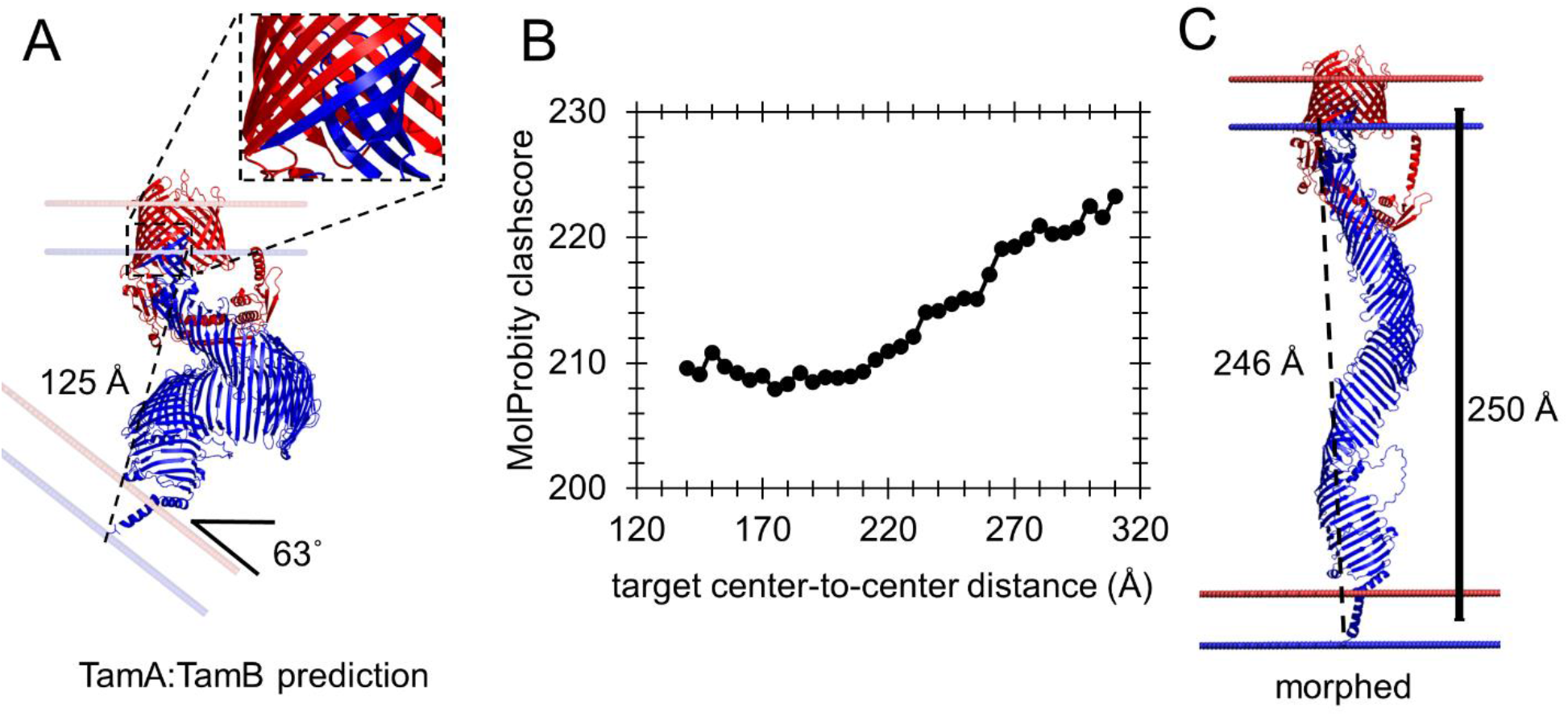
Modeling the complete Tam TEC. (A) The TamA:TamB AF2Complex prediction with the highest piTM score is shown with a zoomed image of the interaction between TamA and TamB (insert). (B) Plot of MolProbity clashscores versus the target center-to-center distances used as input in membrane morphing simulations of the complete Tam TEC. (C) The final structure of the Tam TEC from membrane morphing simulations using a target center-to-center distance from previous estimates of the periplasmic width (see main text).

While the TamA:TamB interface appears to be well modeled in the AF2Complex-predicted structure, the overall shape of the TamB component is highly twisted, and the predicted planes of the IM and OM are tilted at an angle of ∼60° relative to one another (Figure 9A). The contorted shape predicted for the TamB component of the complex is qualitatively consistent with the TamB-only AlphaFold2-predicted structure that is already available (Figure 8C). But the twisting is so pronounced that the entire protein spans only ∼125 Å, which is much shorter than estimates of the width of the periplasm (Graham et al., 1991; Oliver 1996; Volmer & Seligman 2010). We therefore performed membrane morphing simulations covering a wide range of center-to-center IM-OM distances. For all tested distances, a significant unwinding of TamB occurred during the simulations, but despite these large-scale movements, the continuous β-sheet structure of the protein was maintained. While we found that a membrane center-to-center distance of 175 Å produced the lowest MolProbity clashscore (Figure 9B), the characteristic parabolic dependence of the clashscore on the center-to-center distance seen with the other TECs was much less pronounced (Figure 9C). Since this suggests that, for this particular TEC, the membrane morphing protocol produces plausible models but no clear “winner”, we opted to select a center-to-center distance of 250 Å for our most likely final model (Figure 9D). A movie showing the membrane morphing is provided in Movie S8.

The highly twisted nature of the AF2Complex-predicted structures of the Tam complex suggested to us that a high degree of conformational strain might have been erroneously built in to the AlphaFold2-predicted structures of TamB. As one way to explore this idea, we performed an explicit-solvent molecular dynamics (MD) simulation using the AlphaFold2-predicted structure of TamB as a starting point (see Figure 10A; STAR Methods). Strikingly, the protein immediately started to unwind during the simulation, and over the course of a 100 ns trajectory the end-to-end distance rose from ∼180 to ∼250 Å (Figure 10B). As was the case with the membrane morphing simulations, this large-scale, spontaneous change in structure was accomplished without any disruption of the protein’s continuous β-sheet structure. In fact, the structure obtained after 100 ns of MD superimposes (Figure 10C) on to the structure obtained from membrane morphing using a center-to-center distance of 250 Å with a Cα RMSD of ∼7 Å which, for a protein so large, is indicative of a substantial degree of similarity. The MD simulations support, therefore, the notion that the predicted structures of TamB obtained from AlphaFold2 and AF2Complex are subject to a degree of conformational strain, and that the structures predicted by membrane morphing are reasonable. A movie illustrating the 100 ns MD simulation is provided in Movie S9.

**Figure 10.**
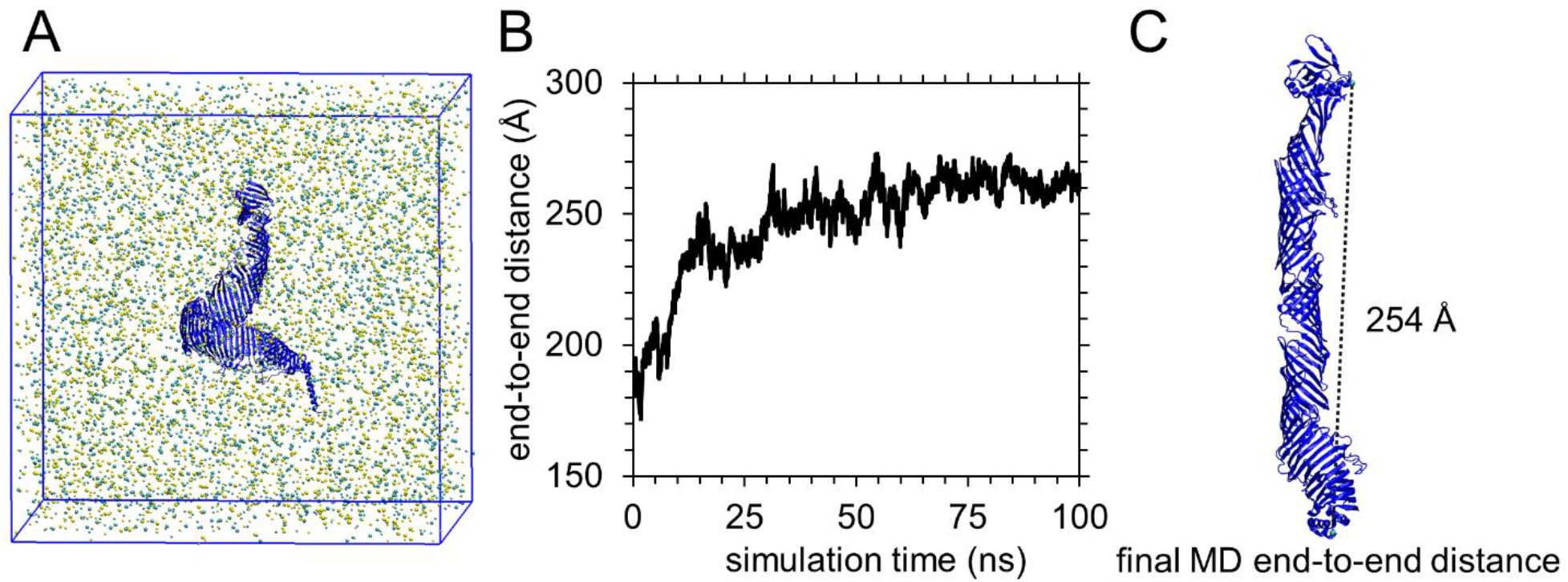
Molecular dynamics simulation of TamB. (A) Image of the TamB simulation system showing the starting TamB AlphaFold2 structure as blue cartoon and ions as spheres colored by name. (B) Plot of the TamB end-to-end distance measured between the Cα atom in the first and last residue versus simulation time. (C) The final TamB structure from the simulation showing the measured end-to-end distance. All structure representations were generated in either PyMol or VMD.

## Discussion

To our knowledge, the structures reported in this work represent the first truly complete atomic models of the principal TECs involved in lipid transport in *Escherichia coli*. At the heart of the approach used here is obviously the combination of DeepMind’s AlphaFold2 method (Jumper et al., 2021) and the Skolnick group’s adaptation of it, AF2Complex (Gao et al., 2022a), which was the first published AlphaFold2-based method focused specifically on the prediction of protein complexes. But the simple membrane morphing protocol reported here also plays a critical role in producing final complex models that are consistent with the conformational restrictions placed on a TEC when it is positioned in its cellular home, spanning the periplasmic space. Importantly, the large distance spanned by a typical TEC means that any structural distortions that result from the membrane morphing simulations can be distributed throughout the entire length of the TEC without needing to be localized to one or a few critical points. The membrane morphing approach depends, in turn, on the Lomize group’s OPM method (Lomize et al., 2022) for predicting the orientations of proteins within membranes. As such, the present study illustrates the potential power of combining a variety of complementary computational techniques – operating within known experimental constraints – for building atomic models of large complexes that might be expensive or very time-consuming to pursue with more conventional experimental methods.

With that said, it is important to note the key assumptions built into the construction of the models presented here. Obviously, the method rests heavily on the assumption that AlphaFold2- derived methods such as AF2Complex can produce accurate predictions of the structures of protein complexes. We think that this has largely been illustrated by the Skolnick group previously (Gao et al., 2022a; Gao et al., 2022b), but an important result obtained here is that – at least for complicated combinations of subunits – repeated predictions may be required to find the most reasonable model. The best example of this is provided by our results for the PqiB_6_:PqiC_6_ interaction, for which the piTM score for one predicted structure was superior to 49 other predicted structures by a large margin (Figure 3F). Crucially, this predicted structure was not only anomalous in terms of its high piTM score, it was also the most biologically plausible of the structures predicted for this complex. At the time of writing, we know additional anecdotal cases where repeated prediction runs have been needed to find a structure that is biologically plausible. It will be interesting to determine, therefore, whether this appears to be a general requirement for complexes with complicated stoichiometries.

Apart from rerunning predictions with different random seeds, we have largely used the AF2Complex method “out of the box”, i.e. without any modifications. As such, we used metrics that are automatically generated by the method as our primary means of identifying the “best” predictions. In general, we found that both the piTM score and the so-called interface score (Gao et al., 2022a; Gao et al., 2022b) performed well, with the former appearing more useful especially when we needed to identify the homo-oligomeric state of a protein, as it seemed less dependent on the number of chains included in the prediction. However, since neither score explicitly monitors nor penalizes the presence of steric clashes, we encountered several cases where predicted structures with promising scores were riddled with clashes due to an absurd placement of identical subunits on top of each other. AlphaFold2’s occasional tendency to overlook obvious steric catastrophes has been noted by others previously (Gao et al., 2022a; David et al., 2022) and has, in part, prompted retraining of the method specifically for complexes by the DeepMind team (Evans et al., 2022); an updated implementation of the AF2Complex method (Gao et al., 2022b), which was published after this work was begun, now uses these retrained networks. While the problem is fortunately easy to identify when it occurs, the fact that it can occur at all indicates that scoring metrics that focus only on a protein-protein interface should be combined with other measures to fully assess the plausibility of a predicted structure.

Aside from our central assumption that AF2Complex is sufficiently accurate that its predictions of the structures of TECs can be believed, there are other key assumptions of our approach that are important to note. One critical assumption that is specific to our models of the Pqi and Let TECs concerns the oligomeric states of PqiA, PqiC, and LetA. We have followed several lines of evidence to conclude that the two IM-resident proteins, PqiA and LetA, are monomers. In both cases, we first examined protein copy numbers estimated from the Weissman group’s ribosome profiling experiments to obtain rough estimates of their likely oligomeric states. This approach is justified on the grounds that relative copy numbers derived from ribosome profiling have previously been shown to correlate well with the known stoichiometries of protein complexes in *E. coli* (Li et al., 2014). PqiB and LetB have both been experimentally demonstrated to be hexamers, and their estimated copy numbers are double those of their binding partners, PqiA and LetA, respectively. This suggests, therefore, that the oligomeric states of PqiA and LetA are likely to be at most trimeric, although we think it is dangerous to attempt to assign the oligomeric state with any higher precision solely on the basis of ribosome profiling.

OPM calculations on the AlphaFold2-predicted structures of the PqiA and LetA monomers produce favorable membrane transfer free energies, thereby suggesting that they are both plausible models for IM-resident proteins. But it was largely on the basis of the following results that we concluded that PqiA and LetA are probably monomeric. First, AF2Complex predictions for homodimeric and homotrimeric forms of both PqiA and LetA provided no compelling evidence of any meaningful inter-monomer interface (Figures 3A and 6A, respectively); this was in marked contrast to what we saw with PqiC (see below). Second, AF2Complex predictions for monomeric PqiA and LetA interacting with their hexameric partners immediately produced plausible structures: in both cases, the PqiA and LetA monomers were contained within what appears to be a loose “cage” formed by the six transmembrane helices contributed by each chain of their hexameric binding partners, and the resulting complexes had very favorable transfer free energies according to OPM. In contrast, while we made repeated predictions for dimeric and trimeric PqiA and LetA interacting with hexameric PqiB and LetB, respectively, their piTM scores did not exceed those found with monomeric variants.

We followed a similar approach in an attempt to assign the oligomeric state of PqiC. The Weissman group’s estimated copy numbers for the known hexamer PqiB and for PqiC suggest that the latter is almost certainly homo-oligomeric, but the exact size of PqiC’s oligomeric state is again difficult to pinpoint further. It is on the basis of the following results, therefore, that we conclude that PqiC is likely also to be a hexamer. First, while AF2Complex predictions for homo-oligomers containing fewer than six PqiC monomers produce the highest piTM scores (Figure 3B), they fail to produce the closed ring structure that we expect given the tube-like shape of its partner PqiB. Second, the internal diameter of the ring structure obtained from AF2Complex predictions of the structure of hexameric PqiC matches nicely with the external diameter of the hexameric PqiB needle. Finally, AF2Complex’s best predicted structure for the PqiB:PqiC complex, while only found in one out of 50 attempts (!), is by far the most plausible structure that we obtained: in it, the PqiB_6_ needle fits snugly within the PqiC_6_ ring, and the N-termini of the PqiC chains – which are known in reality to act as lipoprotein signal peptides – are directly oriented toward the expected position of the OM.

Given the above, we think that there is reasonably good evidence to support our assignments of the oligomeric states of PqiA, PqiC, and LetA. Obviously, however, if our assignment of either PqiA or LetA as monomers ultimately proves to be wrong, then the IM-proximal section of our proposed models of the complete Pqi and Let TECs will need to be extensively revised. Similarly, if our assignment of PqiC as a hexamer ultimately proves to be incorrect, then the OM-proximal section of the complete Pqi TEC model will require revision.

A second central assumption of our approach is that a molecular mechanics-based membrane morphing protocol can be used to make minimal adjustments to TEC models so that they traverse the periplasmic space in an orientation perpendicular to the planes of both the IM and the OM. The extent to which membrane morphing distorts a TEC from its initial predicted structure depends, of course, on: (a) the extent to which the OPM-predicted planes of the IM and OM are tilted with respect to each other in the initial model, and (b) the assumed distance separating the centers of the two membranes (see below). For most of the TECs studied here, the OPM-predicted planes of the IM and OM are already largely parallel to each other, but even when they are drastically tilted such as in the initial model of the TamAB complex (Figure 9A), the membrane morphing protocol acts effectively to bring them into alignment.

The role of the assumed distance separating the centers of the two membranes is potentially more of an issue. For most of the TECs studied here, we determined the optimal center-to-center distance by first using membrane morphing simulations to build a range of models and then selecting as our final model the one producing the lowest MolProbity clashscore (Davis et al., 2007; Chen et al., 2010; Williams et al., 2018). This has the advantage of being an automatable, objective approach, and importantly the optimal models identified using this approach corresponded well with those selected by the much more subjective method of visual inspection. The only TEC for which this approach produced equivocal results was Tam: the highly contorted nature of the AF2Complex-prediction for this complex almost certainly makes it a special case (see below).

For the other TECs, the optimal center-to-center distances identified using MolProbity’s clashscore appear consistent with what is already known about the width of the periplasm. Perhaps the most direct way to obtain an independent estimate of the distance likely to be spanned by an *E. coli* TEC is by applying the same OPM methodology to rigid TECs whose structures have been solved experimentally. To this end, we examined the structure of the AcrABZ-TolC TEC (RCSB ID: 5NG5; Wang et al., 2017) and obtained an OPM-predicted center-to-center distance between the IM and OM of ∼270 Å; doing the same for the structure of the MacAB-TolC complex (RCSB ID: 5NIK; Fitzpatrick et al., 2017) gave a value of ∼275 Å. These values are similar to the optimal distances obtained here for the Pqi and Let complexes: 265 and 240 Å respectively. For the Lpt TEC, the center-to-center distance obtained for the most commonly cited variant (that containing 4 copies of LptA) is 290 Å. It seems likely, however, that this TEC’s extended β-sheet structure would allow it a considerable degree of conformational flexibility that may allow it to accommodate a range of IM-OM center-to-center distances. In the future, it might be interesting to explore the potential flexibility of the Lpt TEC using MD simulations of the kind already used to study other TECs (nicely reviewed in Khalid et al., 2022).

The Tam TEC deserves further discussion. The predicted structure of TamB appears unusually twisted and compact in both its isolated structure (Figure 8C) and in the AF2Complex-predicted structure of its complex with TamA (Figure 9A). It seems quite possible that this compaction is attributable in part to a known artifact of AlphaFold2 that can drive proteins whose structures should be largely extended into roughly spherical arrangements (Ruff & Pappu 2021; Bruley et al., 2022). It is also possible, however, that the compaction results from small errors at each of the strand:strand interfaces that become compounded along the length of the 68-stranded β-sheet structure: it should be remembered, of course, that AF2Complex has no knowledge that a TEC needs to traverse the periplasmic space and so has no reason to explicitly build proteins such as TamB in conformations that ensure their IM- and OM-proximal regions are widely separated.

Regardless of its origins, the highly twisted nature of TamB’s predicted structure suggests the presence of some degree of conformational strain, and this impression appears to be confirmed by the MD simulations that we performed: within 100 ns, TamB spontaneously extends itself to a length sufficient to bridge the periplasmic space (Figure 10B). Interestingly, the structure obtained at the end of the MD simulation is similar to that obtained from membrane morphing using a IM-OM center-to-center distance of 250 Å. The MD simulation required 14 weeks of computer time to conduct on a server equipped with four very capable GPUs (see STAR Methods). Since the membrane morphing simulation of the complete Tam TEC took less than an hour, membrane morphing appears to provide a very rapid strategy for circumventing the need for more traditional simulation approaches to allow conformational relaxation of AlphaFold2 structures. It seems likely, then, that a similar approach will prove useful for building models of other protein complexes that are subject to multiple orientational constraints when placed in their cellular homes.

## STAR Methods

### Summary of overall approach

For each TEC, we started by using the extremely useful SEQATOMS Blast server (Brandt et al., 2008) to identify all available experimental homologous structures for each chain of the complex; when available, all of our predicted models were compared with these structures to ensure their accuracy. Most of the TECs studied here contain total residue counts that exceed the 2400-residue limit recommended by the developers of the underlying AlphaFold2 pipeline (Jumper et al., 2021). For this reason, a divide-and-conquer approach was adopted, when needed, whereby the TEC was split into smaller subcomplexes for which predictions could be more easily carried out. Structure predictions of subcomplexes were performed with AF2Complex (Gao et al., 2022a), and the most plausible predicted structures were combined to build a complete atomic model of each TEC by superimposing residues shared between subcomplexes using TM-align (Zhang & Skolnick, 2005). The models produced in this way were energy minimized in GROMACS (Abraham et al., 2015) to eliminate steric clashes, and then subjected to OPM calculations (Lomize et al., 2022) in order to predict the position and orientation of both the *E. coli* inner and outer membranes. Membrane morphing simulations were then used to make sure that IM- and OM- resident components were properly situated in their respective membranes. Finally, a second energy minimization was performed in GROMACS and the quality of each finalized structure was assessed using the MolProbity webserver (Davis et al., 2007; Chen et al., 2010; Williams et al., 2018). Details of each of these stages are provided below.

### AF2Complex predictions

All structure predictions reported here were performed with the AlphaFold2-derived method AF2Complex (Gao et al., 2022a). The AF2Complex submission script automatically reports a variety of metrics that can be used to assess the plausibility of a predicted structure. In general, we found the piTM score to be the most informative of these measures. While pLDDT scores were less useful in our hands for assessing the quality of interactions, they were sometimes helpful for flagging problems in predicted structures that were not apparent from their piTM or interface scores. In particular, in those pathological cases where the AlphaFold2 engine placed copies of a protein directly on top of each other, the pLDDT score often dropped significantly.

In most literature uses of AlphaFold2 and its derived methods, it is common to run the prediction only once: the submission script automatically generates one predicted structure from each of the five neural network models trained by the DeepMind team. The submission script, however, provides for the possibility of setting an initial random seed and we found that, for some of the more complicated cases examined here, quite different results could be obtained if this random seed was changed. For certain subcomplexes, therefore, we performed ten independent submissions, with the total number of predicted structures obtained in such cases amounting to 50. Our strategy for deciding which subcomplexes to treat in this way was somewhat subjective. In general, we started by examining the five piTM scores produced from the first submission of the AF2Complex script. If the best of these piTM scores was close to, or above, the 0.50 threshold recommended by the Skolnick group *and* if the resulting structure looked plausible, then we carried out no further predictions. If, however, the best structure produced only a poor piTM score or was clearly implausible, then we made a further nine submissions of the script in the hope of obtaining a better result.

All predictions reported here were performed on the University of Iowa’s High Performance Computing Center using machines equipped with either NVIDIA GEForce RTX 2080Ti or NVIDIA A10 GPUs. The runtimes and the RAM requirements for predictions varied significantly, but typical predictions were completed within a day while the longest predictions required up to a week.

### Construction of initial models of complete TECs

Once plausible structural models of subcomplexes were obtained, they were combined to build initial models. This was achieved using superpositions performed with TM-align (Zhang and Skolnick, 2005). The choice of which set of superimposed coordinates to retain in the complete model was made on a case-by-case basis, but preference was always given to experimentally determined coordinates when they were available. While the superposition of fragments of subcomplexes generally worked well for each of the TECs examined here, steric clashes between elements of structure not involved in the superpositions did occasionally occur. In an attempt to mitigate the effects of any such clashes, energy minimizations were performed with GROMACS (see below).

### OPM calculations

In order to determine how each of the complete TEC models might be situated in the two membranes, we submitted each of the energy-minimized models to an in-house installation of OPM v3.0 (Lomize et al., 2022). These predictions were performed using two sets of input parameters: one for aligning the position of the IM and another for aligning with the OM. For each complex we used the input parameter file type "2" that is included with the source code and that allowed us to specify: (a) that the prediction should use planar Gram-negative membrane types of "GnI" and "GnO" for the IM and OM, respectively, and (b) the identities of the chains to be inserted in each membrane. Following prediction, OPM returns a (z-axis-aligned) membrane-oriented version of the input .pdb file centered on the origin of the coordinate system, and adds in HETATM entries whose z-coordinates define the upper and lower limits of the predicted membrane; the latter are used to identify those atoms that are membrane-embedded (see below).

### Membrane Morphing Protocol

The membrane morphing simulations devised here make use of in-house code that takes as input two initial structures oriented and anchored in the IM and OM by OPM and performs a series of GROMACS energy minimizations that drive the OPM-predicted center of the OM to a user-specified distance from the IM. The code first reads the IM-anchored .pdb file and identifies those atoms that should remain embedded in the IM by comparing their z-coordinates with the minimum and maximum z-coordinates assigned to the IM by OPM. The atomic coordinates read from the IM-anchored .pdb file serve as the initial coordinates for the subsequent membrane morphing simulations (see below). The code then reads an OM-anchored version of the same .pdb file (also predicted by OPM) and identifies those atoms that are to remain embedded in the OM by comparing their z-coordinates with the minimum and maximum z-coordinates assigned to the OM by OPM. For obvious reasons, atoms cannot be assigned to be embedded in both the IM and the OM.

The code then prepares for membrane morphing simulations by constructing a generic molecular mechanics energy model that is intended to retain the structure read from the input IM-anchored .pdb file. Importantly, since the energy model contains only “bonded” interactions of the type that are routinely used in molecular mechanics applications, it contains no electrostatic or Lennard-Jones terms. The energy model is constructed in the following way. Conventional covalent bonds are first added between: (a) atom pairs within the same residue that satisfy simple distance-based criteria, and (b) atom pairs that form peptide bonds (i.e. atom “C” of residue *i* and atom “N” of residue *i+1*). Harmonic potential functions (GROMACS bond type “1”) are used to restrain these bonds at their initial values with a force constant of 41840 kJ/mol/nm^2^. Conventional angle terms are then added between all bond pairs that share a common atom, and harmonic potential functions (GROMACS angle type “1”) are used to restrain them at their initial values with a force constant of 83.68 kJ/mol/rad^2^. Conventional dihedral terms are then added between all angle pairs that share two bonded atoms, and single-cosine potential functions (GROMACS dihedral type “9”) are used to restrain them at their initial values with: (a) a barrier half-height of 4.184 kJ/mol for dihedrals in which the two central atoms are both sp^3^-hybridized, or (b) a half-height of 83.68 kJ/mol for dihedrals in which one or both of the central atoms are sp^2^-hybridized. The much higher energetic barrier assigned to rotation of dihedrals that involve sp^2^-hybridized atoms helps to ensure that the planarity of peptide bonds and aromatic groups is maintained during membrane morphing. Finally, to maintain the tertiary and quaternary structure during morphing, additional harmonic potential functions (GROMACS bond type “6”) are added between all nonbonded atom pairs that are within 6 Å in the IM-anchored initial .pdb file; as with the conventional bonds, these additional “bonds” are assigned a force constant of 41840 kJ/mol/nm^2^.

Having defined an energy function that seeks to maintain the initial structure read from the IM-anchored .pdb file, the only additional terms included in the simulations are flat-bottomed position restraints (of GROMACS type “2”) that are applied to membrane-embedded atoms to restrain them to a user-specified reference z-coordinate. These position restraints are applied to all atoms that are identified as being embedded in either the IM or the OM (see above); during simulations, the restraints act to maintain these atoms at the appropriate depth within their respective membranes. While the position restraints are formally of the “flat-bottomed” type, in practice we assign no “flat” region to them: instead, the restraint function is a simple harmonic potential function with a force constant of 10000 kJ/mol/nm.

The code then conducts membrane morphing simulations in the following way. A series of energy minimizations is performed during which the reference z-coordinates used to restrain the OM-embedded atoms are linearly interpolated from their initial values to their final desired values. These initial values are taken directly from the atom’s z-coordinates in the IM-embedded .pdb file provided by the user; their final values are taken from the atom’s z-coordinates in the OM-embedded .pdb file but incremented by the desired final center-to-center distance between the two membranes. The reference z-coordinates for the IM-embedded atoms remain unchanged throughout the entire process. For all of the TECs discussed here, the interpolation of the OM-embedded atom z-coordinates was carried out in 100 stages, and an energy minimization was performed using GROMACS at each stage; the structure obtained at the end of each stage was then used as the initial structure for the next stage of the protocol. Each energy minimization consisted of a maximum of 1000 steps using the conjugate-gradient algorithm with an initial stepsize (`emstep`) of 0.01 nm; early termination of the energy minimization could occur if the energy changed by less than a specified tolerance (`emtol`) of 0.1 kJ/mol.

### Energy minimization of atomic structures using GROMACS

While the membrane morphing protocol involves energy minimizations of a purely bonded energy model, more conventional GROMACS energy minimizations were also used to eliminate steric clashes from our atomic structures. Such energy minimizations were carried out at two stages of the TEC building process: (a) immediately after superimposing subcomplexes to produce initial models of complete TECs, and (b) immediately after the final stage of membrane morphing. The energy model used in all of these minimizations was the CHARMM36m force field (Huang et al., 2017): a 12 Å cutoff was applied to all nonbonded interactions, and electrostatic interactions were calculated using a relative dielectric constant of 78.4 in order to crudely mimic the screening effects of water molecules since these were not explicitly modeled. These energy minimizations were conducted within GROMACS (Abraham et al., 2015) using the steepest descent algorithm for 1000 steps with an initial step size (`emstep`) of 0.0001 nm.

### Assessment of structure quality using MolProbity

The quality of each morphed structure was assessed using the MolProbity webserver (Davis et al., 2007; Chen et al., 2010; Williams et al., 2018) with default options. Each structure was uploaded to the webserver, and the “Analyze geometry without all-atom contacts” tab was selected. The following options were then selected: under the “3-D kinemage graphics – Universal” subsection we selected “Clashes”, “Hydrogen bonds” and “van der Waals contacts”; under the “Charts, plots, and tables – Universal” subsection we selected “Clashes & clashscore”. While many different metrics are reported by the webserver, our focus was on the reported “Clashscore, all atoms” which is defined as “the number of serious steric overlaps (> 0.4 Å) per 1000 atoms.”

### Molecular dynamics simulation of TamB

An explicit-solvent MD simulation of TamB was performed starting from the AlphaFold2 structure already available in the AlphaFold2 database (https://alphafold.ebi.ac.uk/files/AF-P39321-F1-model_v4.pdb) using GROMACS 2020 (Abraham et al., 2015) with the CHARMM36m forcefield (Huang et al., 2017). Hydrogens were first added with the GROMACS utility `pdb2gmx`; the structure was then placed in a cubic box using `editconf` with 350-Å sides, solvated using `solvate` with the CHARMM-TIP3P water model (Jorgensen et al., 1983), and neutralized via `genion` with 150 mM NaCl. The system was energy minimized using the steepest descent algorithm and progressively heated in 50-K increments to 298 K over the course of seven 60-ps equilibration stages using the Berendsen thermostat and isotropic barostat (Berendsen et al., 1984). All stages of the simulation used a timestep of 2 fs. Prior to production, the system was equilibrated for an additional 1 ns using the Parinello-Rahman isotropic barostat (Parinello & Rahman, 1981) and Nose-Hoover thermostat (Hoover et al., 1985). The short-range electrostatic and van der Waals cutoffs were set to 12 Å and implemented with the Verlet cutoff-scheme (Páll & Hess, 2013); long-range electrostatics were calculated using the smooth particle mesh Ewald method (Essmann et al., 1995). Covalent bonds involving hydrogen atoms were constrained using the LINCS (Hess et al., 1997) algorithm. A production simulation of the entire system – which comprised 4,206,29 atoms – was run for 100 ns with snapshots saved at intervals of 50 ps. Running on a new server equipped with four NVIDIA A10 GPUs, the simulation required approximately 14 weeks to complete.

## Figure generation

Bar charts, histograms, and scatterplots were made in the statistical programming language R with the ggplot2 package. All schematic figure images were generated using BioRender, while all other images were generated using either PyMol 1.8.4 (Schrodinger, LLC 2015) or VMD 1.9.4 (Humphrey et al., 1996).

## Acknowledgments

This research was supported by a grant from the National Institutes of Health (R35 GM122466) to AHE and supported in part through computational resources provided by The University of Iowa.

## Author Contributions (CRediT Statement)

Robert T. McDonnell: Methodology, Formal analysis, Investigation, Data Curation, Writing – original draft preparation, Visualization. Nikhil Patel: Investigation, Data Curation.

Zachary J. Wehrspan: Methodology, Investigation, Data Curation.

Adrian H. Elcock: Conceptualization, Methodology, Software, Writing – review and editing, Supervision, Project Administration, Funding acquisition.

## Data availability

All finalized TEC structures and all computer code necessary to run the membrane morphing simulations described here will be made available to reviewers at the time of manuscript review. Upon acceptance of the manuscript for publication, all finalized TEC structures will be available as supplementary material, and the computer code will be available to the community at the following GitHub repository (https://github.com/Elcock-Lab/membrane_morphing).

