## Supplemental Figures for "Atomic Models of All Major Trans-Envelope Complexes Involved in Lipid Trafficking in *Escherichia Coli* Constructed Using a Combination of AlphaFold2, AF2Complex, and Membrane Morphing Simulations"

### Slide 1
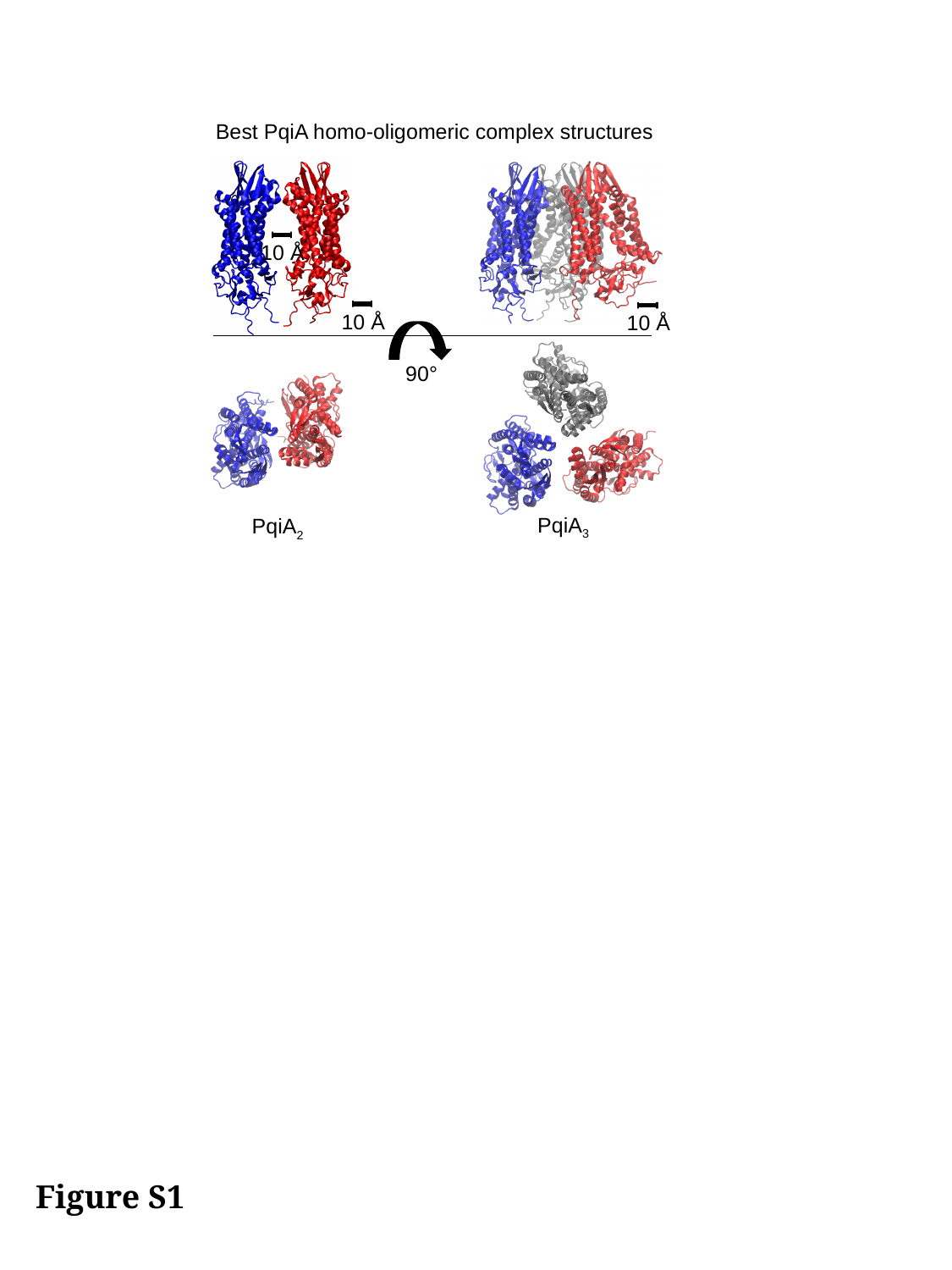

Best PqiA homo-oligomeric complex structures
10 Å
10 Å
10 Å
90°
PqiA_3_
PqiA_2_
Figure S1

### Slide 2
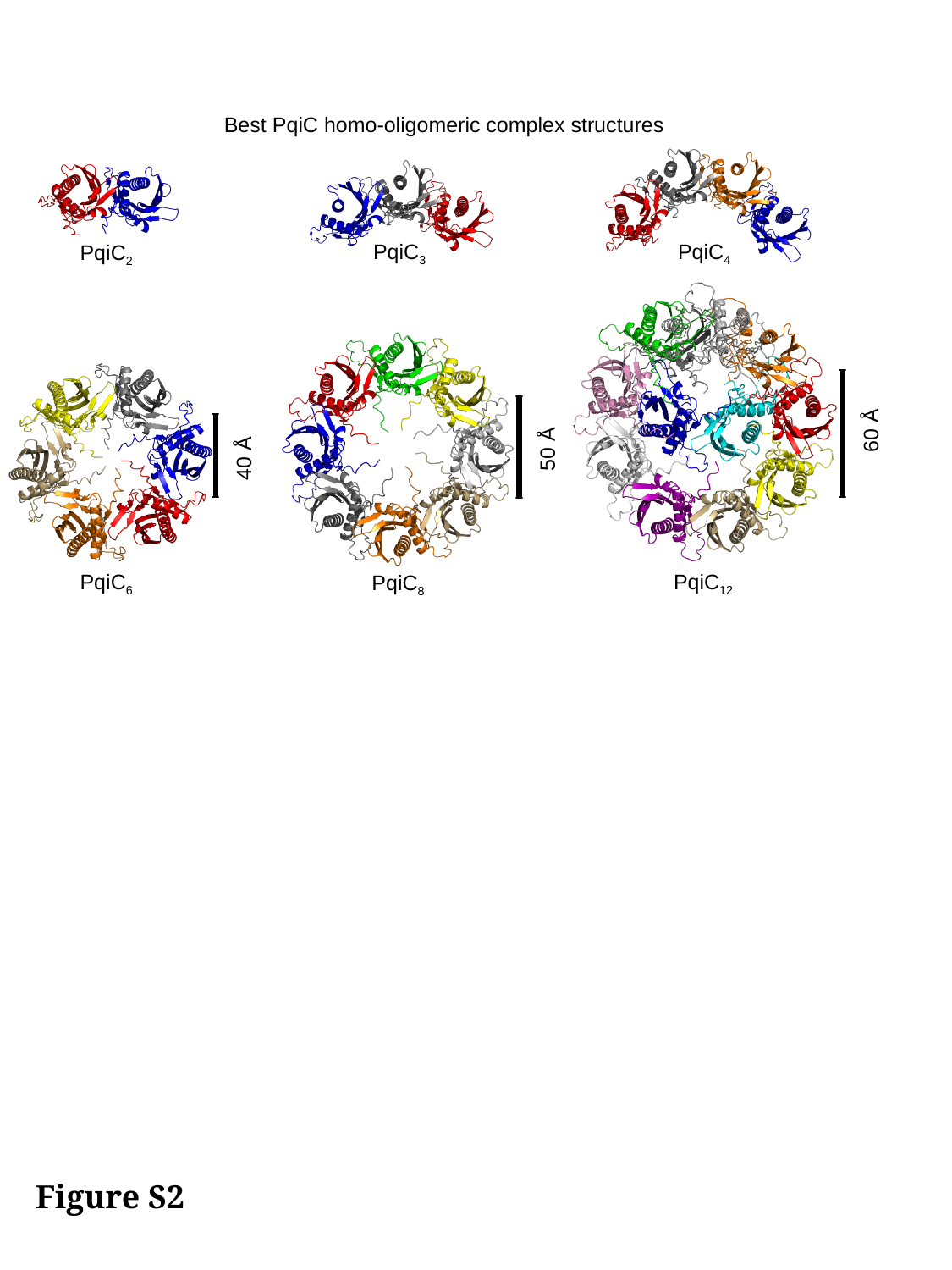

Best PqiC homo-oligomeric complex structures
PqiC_4_
PqiC_2_
PqiC_3_
60 Å
PqiC_12_
50 Å
PqiC_8_
40 Å
PqiC_6_
Figure S2

### Slide 3
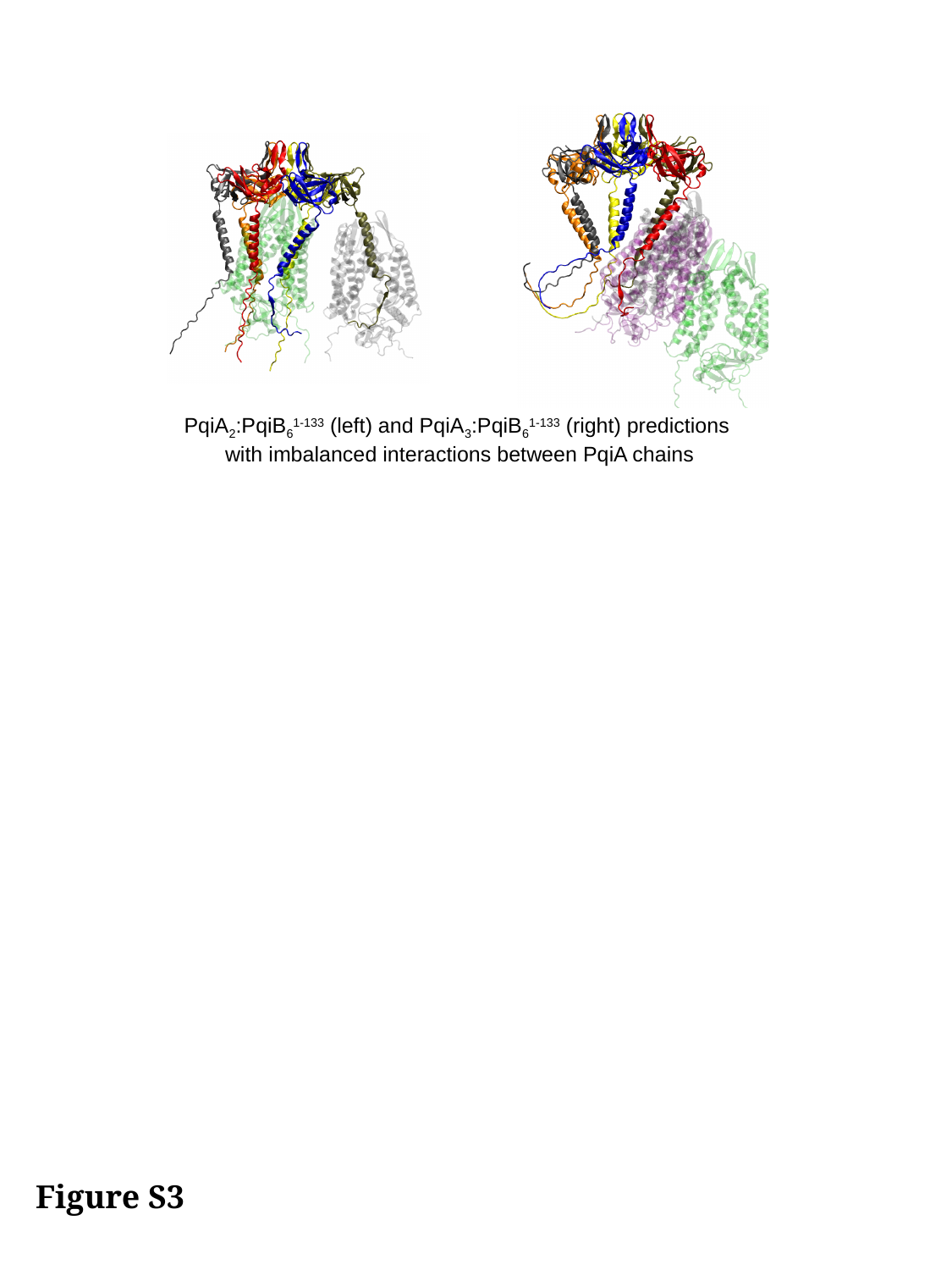

PqiA_2_:PqiB_6_^1-133^ (left) and PqiA_3_:PqiB_6_^1-133^ (right) predictions
with imbalanced interactions between PqiA chains
Figure S3

### Slide 4
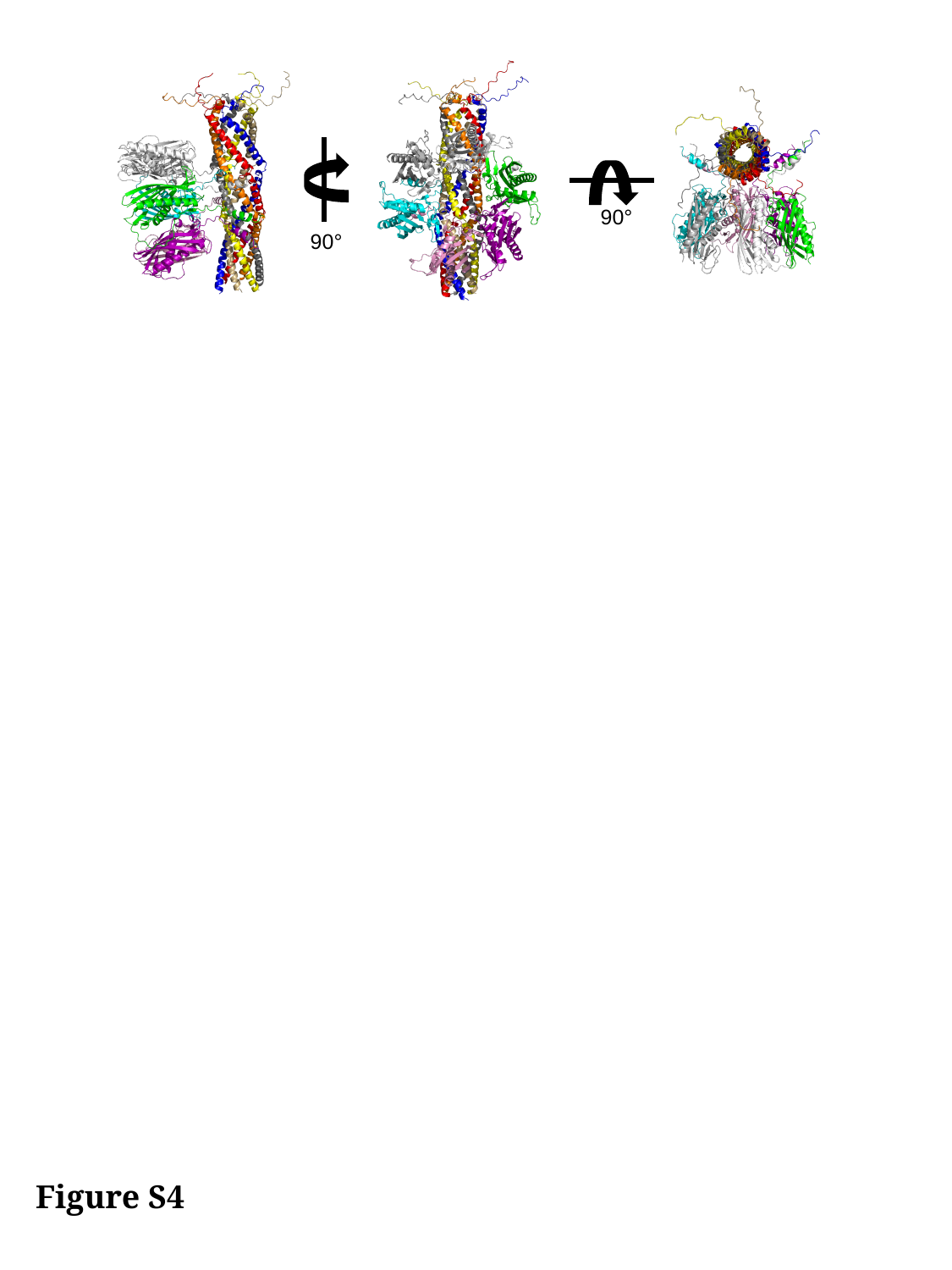

90°
90°
Figure S4

### Slide 5
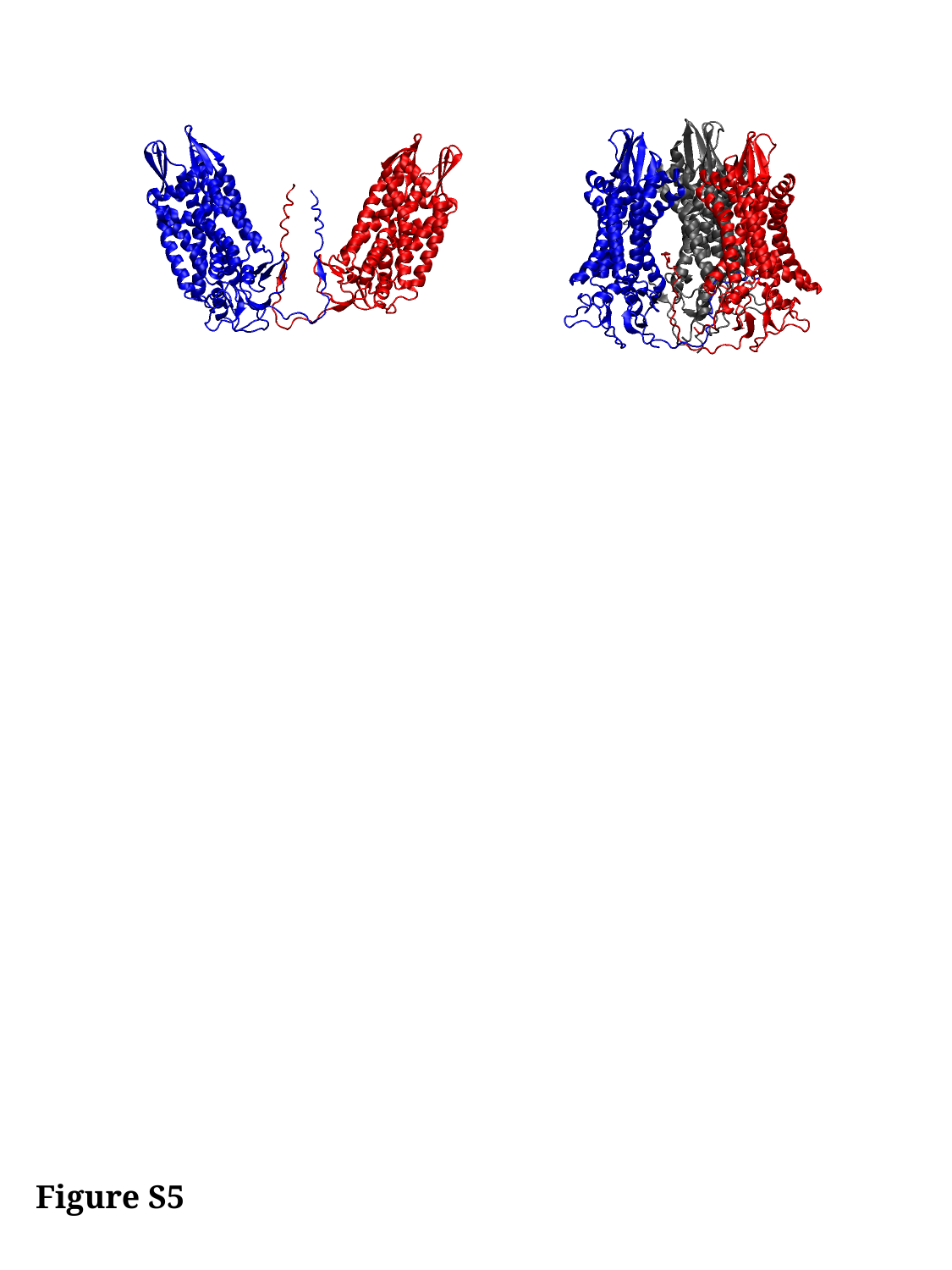

Figure S5

### Slide 6
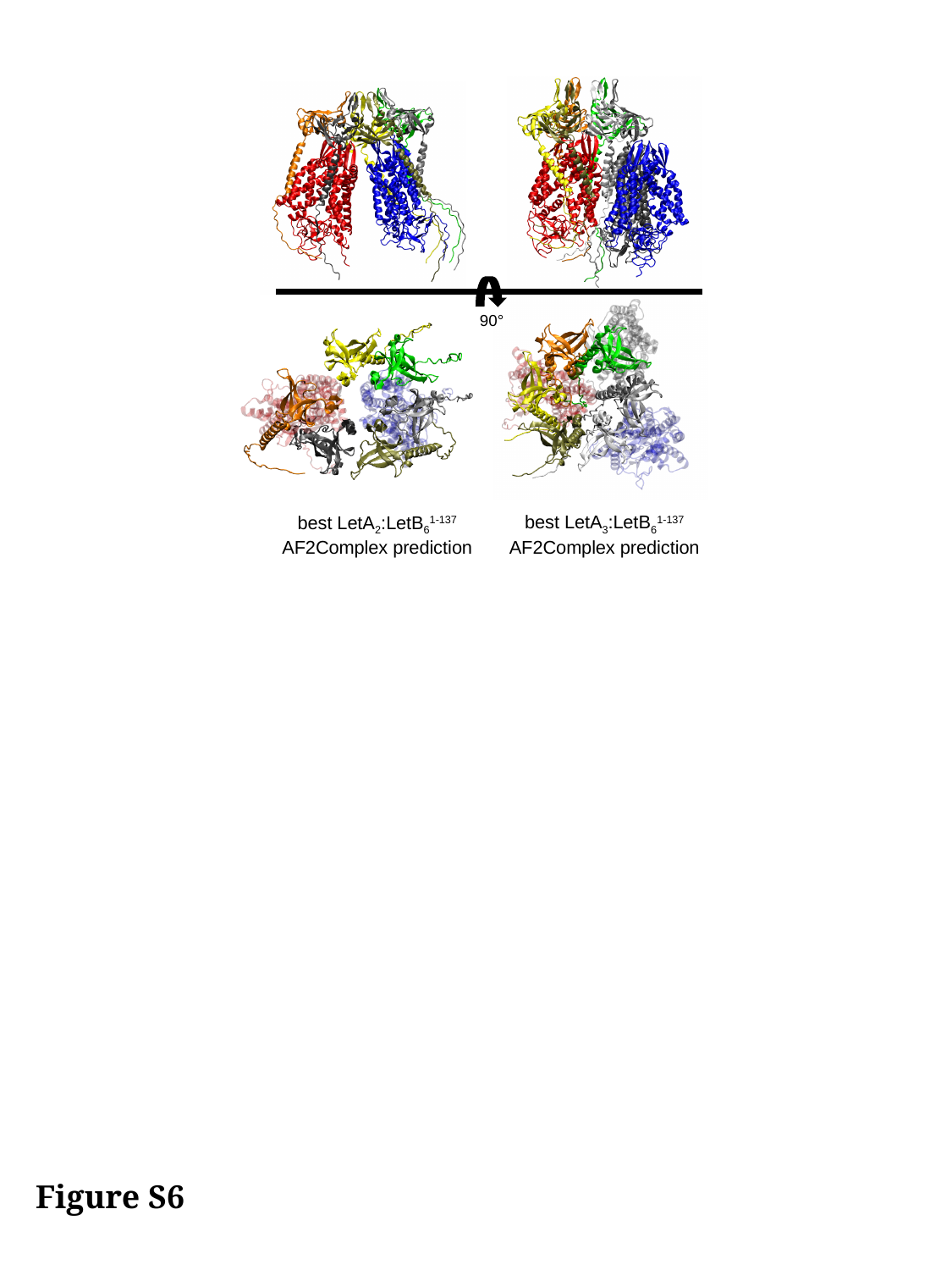

90°
best LetA_3_:LetB_6_^1-137^
AF2Complex prediction
best LetA_2_:LetB_6_^1-137^
AF2Complex prediction
Figure S6
